# Inferring stability and persistence in the vaginal microbiome: A stochastic model of ecological dynamics

**DOI:** 10.1101/2024.03.02.581600

**Authors:** José M. Ponciano, Juan P. Gómez, Jacques Ravel, Larry J. Forney

**Affiliations:** Department of Biology, University of Florida, Gainesville, FL; Departamento de Química y Biología, Universidad del Norte, Barranquilla, Colombia; Institute for Genome Sciences, Department of Microbiology and Immunology, University of Maryland School of Medicine, Baltimore, MD; Institute for Bioinformatics and Evolutionary Studies, Department of Biological Sciences, University of Idaho, Moscow, ID

## Abstract

The vaginal microbiome is dynamic, yet the ecological and stochastic forces shaping its stability and persistence remain poorly understood. We developed a multi-species stochastic population model to analyze time-series data from 135 individuals sampled daily over 70 days, integrating ecological theory with microbial dynamics. Our framework explicitly incorporates stochasticity, ecological feedback, and sampling error to quantify community stability. We show that intra- and inter-species interactions and environmental fluctuations critically shape microbial population trajectories. This approach enabled the estimation of species competition coefficients and the identification of distinct stability regimes across individuals. We also introduce a Risk Prediction Monitoring (RPM) tool to track persistence probabilities of key taxa, particularly *Lactobacillus* spp., mirroring extinction risk models in conservation biology. Our findings challenge static “healthy” vs. “dysbiotic” categorizations and offer a quantitative framework for assessing microbiome resilience. This has direct implications for microbiome-targeted therapies aimed at promoting ecological stability in vaginal bacterial communities.

## Introduction

Inferring the interplay between stochastic processes and the ecological and evolutionary conditions that permit the establishment and persistence of host-associated microbial communities has remained a topic laden with controversies and unresolved conceptual and practical issues ^1–5^. The paucity of studies that connect extensive time-series data with population dynamics models rooted in ecological principles has been at the center of the problems faced when inferring processes from patterns in this area of research. However, even a basic understanding of the mechanisms leading to these fluctuations remains elusive. Given that the structure and composition of an ecological community often alternate between distinct, widely different states^6–8^, the chances of dramatic community shifts are better predicted using mechanistic, stochastic population dynamics models ^9–13^. Illustrating the conceptual and practical advantages of fitting stochastic population dynamics models to multi-species bacterial time series data is the focus of this paper, exemplified here for the human vaginal microbiome.

Indigenous bacterial populations in and on the human body constitute the first line of defense against infection by preventing non-indigenous organisms from causing disease. Considerable efforts have been made to characterize the composition of vaginal bacterial communities found in healthy reproductive-age women and to understand interruptions to the homeostasis of this microbiome. Community compositions that widely differ from these “normal” states are thought to be states of ‘dysbiosis’. Dysbiosis can reflect changes in the absolute numbers of microbes, the species composition, or the relative abundances of bacterial taxa or some combination thereof. Such states reflect an ‘imbalance’ in the vaginal microbiome and are usually considered as ‘unhealthy’ states. For example, microbiome states depleted of *Lactobacillus* species are said to reflect ‘dysbiosis’ despite persisting for extended periods of time in women who are asymptomatic and otherwise healthy. In other cases, it has been recognized that bacterial communities of reproductive-age women often vary over time in a seemingly haphazard way or reflect idiosyncratic changes in species composition^14^, making it challenging to associate states with particular conditions. Nonetheless, temporal fluctuations in the relative abundances of the different species are undeniably associated with specific environmental variables like pH. Understanding these fluctuations that lack pattern can help identify interesting mechanisms associated with health issues and potentially chronic conditions.

Combining mathematical, statistical, and stochastic process tools to explicitly model the mechanisms that underlie community dynamics on a temporal scale has long proved to be a fruitful approach to filling knowledge gaps regarding the functioning of ecological communities ^10,15^. This approach has also been shown to reliably reproduce the regular waxing and waning of natural population densities in single and multi-species systems ^11,12,16–21^. Mathematical characterizations of how the mean and variance of population sizes change over time can be obtained by formulating multi-species population growth as stochastic processes ^4,11,22^. These mathematical expressions reveal the links between the patterns of population variation, environmental variation, and key ecological quantities like intrinsic growth rates and inter-specific and intra-specific competition coefficients.

Stochastic models of the temporal fluctuations of species’ abundances aim to translate fundamental concepts in ecology and evolution into testable hypotheses and predictions that can be confronted with abundance time series datasets. These models decompose the changes in abundances of one, two or more species over time into four main components ^4,10–12,23,24^: 1) basic demographic processes like reproduction and the effects of density dependence and inter-specific interactions, 2) chance variation and individual heterogeneities affecting births and deaths, known as “demographic stochasticity” effects; 3) environmental stochasticity or temporal variation in vital rates (e.g., birth and death rates) that reflect variation in environmental conditions; and 4) observation error and sampling noise. This last component is particularly relevant in microbial systems ^2,4,19,25^.

A practical application of these stochastic population dynamics models was presented by Ives et al ^26^ almost 21 years ago. In the MAR (Multivariate Autoregressive) model, the overall effect of environmental fluctuations in the growth of a population is modulated by ecological processes^26^. These authors showed that the growth rate of a population characterized by weak density-dependence was easily affected by fluctuations in the quality of the environment. In contrast, those populations characterized by strong density-dependence were not. When presented with the same temporal regime of environmental variation, a population with strong density-dependence will fluctuate much less than a population with weak density-dependence. Furthermore, stability could be measured and conceptualized for a single population as the ratio of the magnitude of environmental variation to the strength of density dependence^26^. This finding allows for a direct comparison of the reactions of two different populations to the same environmental noise regime (see Figure 1).

**Figure 1.**
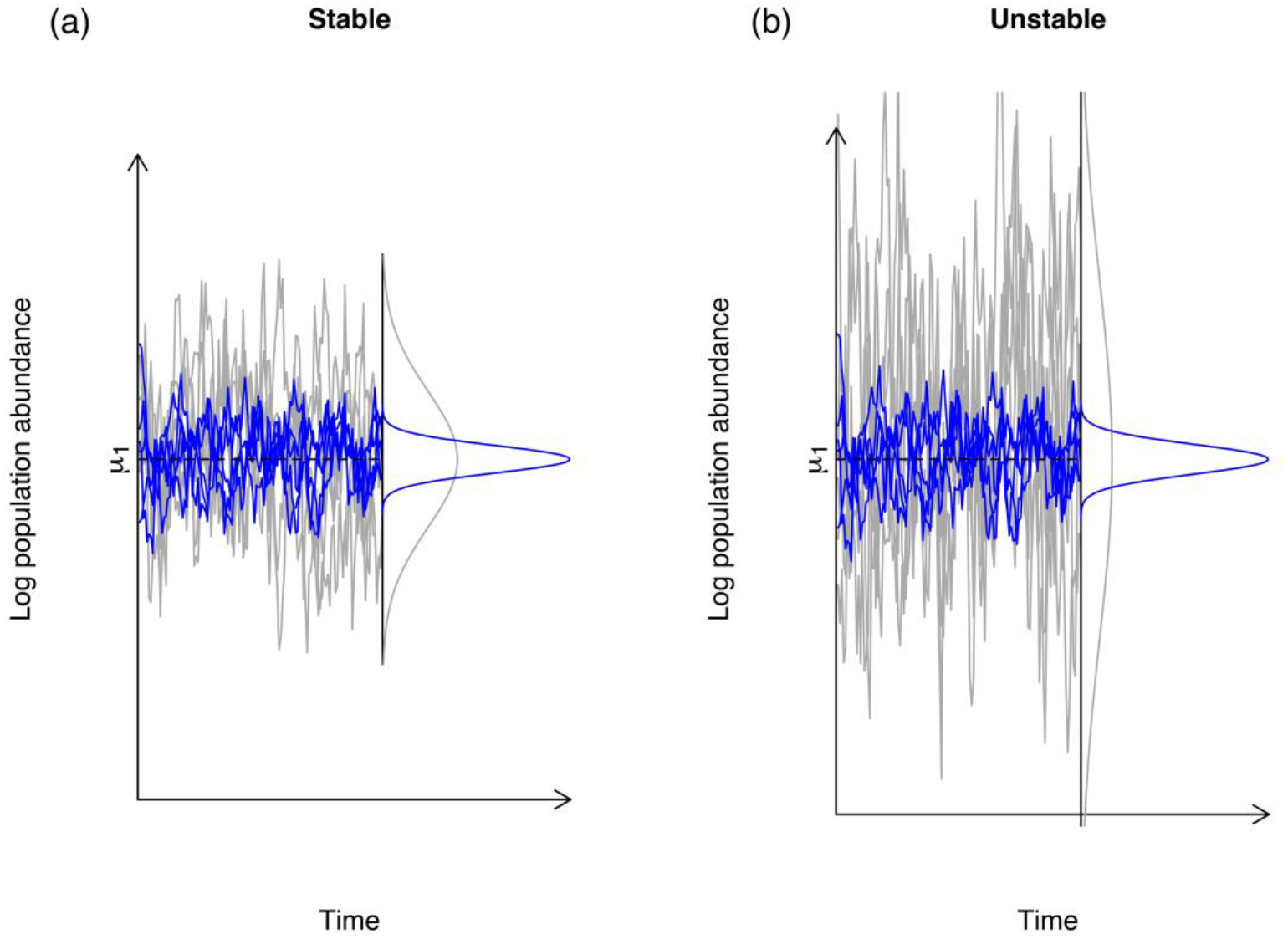
The abundances in stable (a) vs. unstable (b) populations. In both panels, the grey lines representing the log-population abundances at stationarity were simulated under the stochastic Gompertz model of Ponciano et al. (2018) under the same environmental noise regime that are shown in blue. The variance of the long-run log-population abundances is equal to the ratio of the environmental noise variance (here 0.11) to one minus the squared strength of density-dependence c. This coefficient is stronger on the left than on the right. On the left c = 0.75 and so the log-population size variance was 0.11/(1 − 0.75^2^) = 0.2514. On the right panel, density dependence is much weaker, with a coefficient equal to 0.93. (Coefficients closer to 1 are close to density-independence.) The variance of the population abundances under the same environmental noise variance is approximately three times higher: 0.11/(1 − 0.93^2^) = 0.8142. The magnitude of the response of a population to environmental noise, in terms of variability, is modulated by c.

This insight, which was brought about by Ives et al in the context of community ecology^26^, made it possible to compare different populations and communities on the same level playing field (see Figure 2).

**Figure 2.**
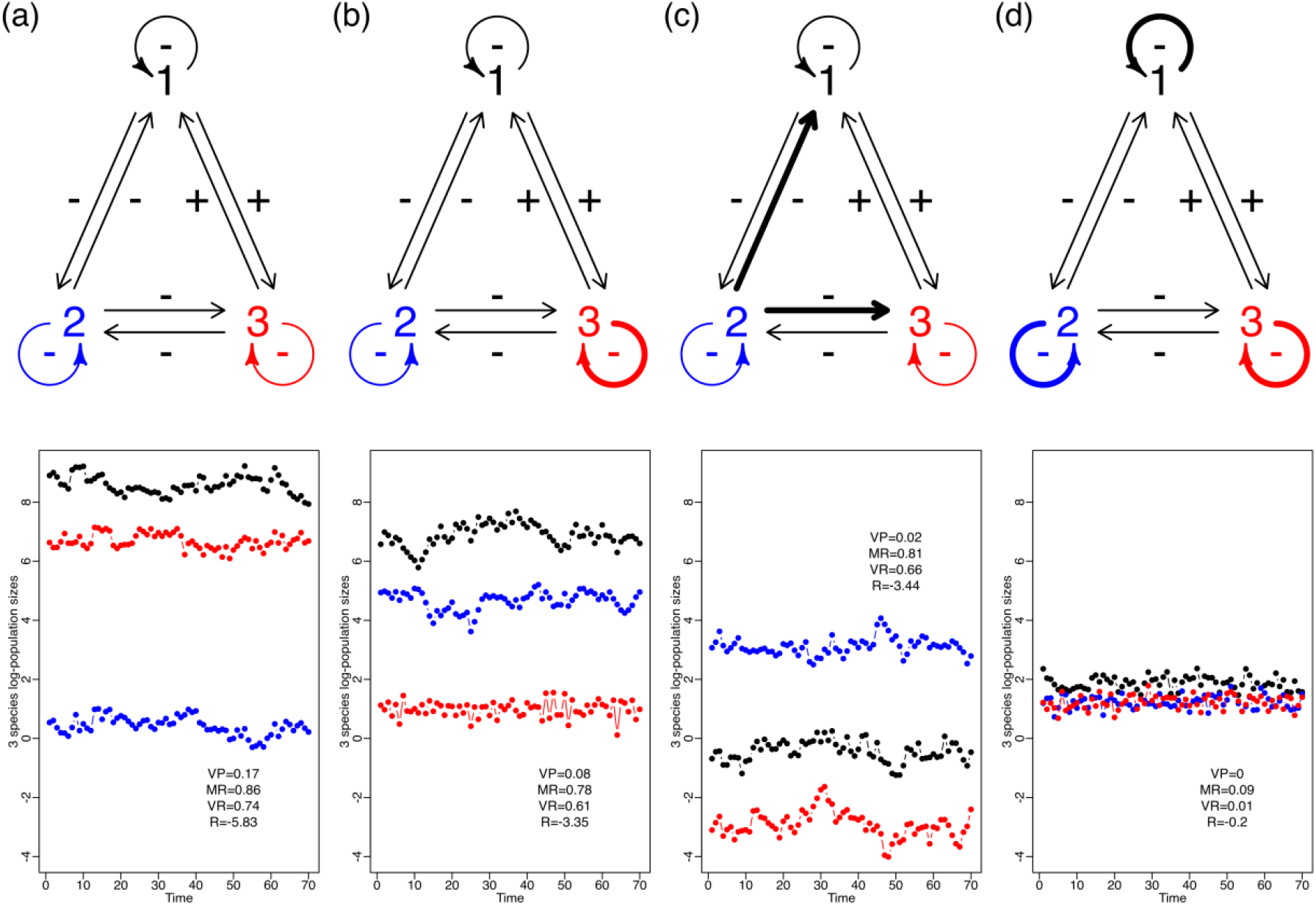
Extending the simulation shown in Figure 1 to two or more species for 70 days. It shows that changes in the nature and intensity of community interactions directly affect stochastic stability, measured using Ives’ *et al*. four stability metrics: VP, MR, VR, and R, which are defined and explained in the main text. In this case, the modulation of a variable environment depends on the structure and the nature and intensity of the intraspecific and interspecific interactions. This figure shows the fluctuation in population sizes of four different community structures (a-d) with three species subject to the same environmental noise regime. The upper row represents the four community structure types. The intraspecific interactions (looped arrows) and interspecific interactions (straight arrows) change in magnitude, with weak interactions shown as thin arrows and strong interactions shown as thick arrows. In the row directly below each of these interaction graphs, we show the resulting temporal dynamics of the log-abundances of each species. Since all four simulations were done under the same environmental noise regime, the differences in magnitude and fluctuation of population abundances across community types can be directly attributed to differences in structure. Weaker interaction strengths (fewer arrows in bold) lead to larger population fluctuations under the same environmental variance. Note that all four Y-axes were made equal to allow for comparison.

In an ecological community, the influence of environmental noise variance is modulated by the density dependence and the inter-specific interaction coefficients ^10,24^. This is illustrated in Figure 2.

Here, we show four scenarios in which intra-specific and inter-specific interactions’ strength varied while environmental noise remained constant. In each of these scenarios, three species (1, 2, and 3) interact in the following ways: species 1 and 2 and 2 and 3 are competitors and thus have a negative effect on each other. Species 1 and 3 are mutualists and have a positive effect on each other (Figure 1). Finally, all the species show negative density dependence. In the first scenario, all the interactions, including intraspecific density dependence, are weak. In the second scenario, only species 3 had strong density dependence, while the rest of the interactions were weak. In the third scenario, species 2 has a strong negative effect over 1 and 3, but the rest of the interactions were kept weak. Finally, we made intraspecific interactions strong in the fourth scenario while keeping interspecific interactions weak. The coefficients used for each scenario are shown in Supplementary Table 1.

We show how the same amount of environmental variance may result in large or small growth rate variation, depending on the maximum growth rates and those specified interaction strengths (Figure 2). In a community, the strength of the inter-specific and intra-specific interactions, and the overall architecture of its assembly ultimately modulates the response to environmental variation. Like in single-species population dynamics, the same level of the observed variation in the growth rate can result from the populations in a community overreacting to mild exogenous fluctuations or, alternatively, from a community dampening considerably unusually large environmental variability. Ives et al showed that it was possible, through the analysis of multi-species time series, to estimate four different statistics or “stability metrics” (called VP, MR, VR and R in Figure 2 (and as explained below) that would allow the comparison of multiple communities in the face of the same magnitude of environmental noise. In essence, and without entering into mathematical details, these authors showed that it was possible through these metrics to obtain a standardized measure of the reaction of a community to such noise. These findings also imply that deeming a particular set of time series as representative of “stable” or “unstable” dynamics just by its overall variability might be misleading and conflate the fundamental processes governing the dynamics of an ensemble of interacting populations.

Here we leveraged the MAR model to develop and test a multi-species stochastic population modeling approach ^11,12,19,22,23,27^ to better understand how fluctuations in the environment ultimately contribute to changes in species composition and abundances as well as to the overall community stability of human vaginal microbiomes along the menstrual cycle of reproductive age women. Our central hypothesis is that stability properties, diversity, and fluctuation regimes of human vaginal microbial communities can be better understood by explicitly accommodating the following three sources of variability in time series models of multi-species abundances: 1) stochastic biotic and abiotic forces, 2) ecological feedback and 3) sampling error. This modeling framework translates tentative explanations of the sources of the temporal variation in bacterial abundances into testable hypotheses that describe the interplay between ecological processes and the dynamics of abiotic factors while considering sampling variability. This translation was achieved by combining time series data of bacterial species composition with stochastic models derived from basic ecological principles. This probabilistic approach results in a practical statistical connection between biological hypotheses and time series data ^12^. Here we exemplify this process using 135-time series of human vaginal microbial communities ^14^.

## Results

### Model formulation

The MAR model is a discrete-time Markov process deeply rooted in stochastic population dynamics modeling theory ^10^. It jointly models three processes that determine the variation in abundance of the species in a community through time: 1) a deterministic density-dependent population growth for every species in the system on a log-scale, 2) the effect of every species on the growth rate of any other species and 3) the effects of environmental variation on the growth rate. This stochastic model has as its deterministic counterpart the multispecies Gompertz density-dependent model, which has been widely applied to estimate bacterial growth^24^. The MAR model is amenable to simulations via recursion because the total abundance of any species in one time step only depends on the abundance of all the species in the previous time step. Thus, the time series data can be modeled using its linear, multivariate recursion and representation.

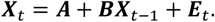

***X***_*t*_ is a vector of the log-population abundances at time *t*, ***A*** is a vector whose elements give the intrinsic rate of increase for each species in the system, ***B*** is a squared matrix whose elements *b*_*ij*_ denote the effect of species abundance *j* on the growth rate of species *i*. Finally, ***E***_*t*_ represents a vector of stochastic environmental factors varying independently from one time step to the next. These factors are modeled with a multivariate normal distribution with mean 0 and variance-covariance matrix **Σ**. Through this variance-covariance matrix **Σ** the modeler can specify whether the response to environmental variation is independent from one species to the next or not, and if not, any covariance structure could be added. In macro-ecological communities, for instance, the response to the environment from one species to the next might be phylogenetically constrained. In bacterial communities, these phylogenetic constraints likely directly translate into explicit functional constraints since any given strain might be better at doing something the others cannot do. The MAR model can be viewed as a linear, first approximation to a complex, multi-species population dynamics process of the form ***n***_*t*_ = *h*(***n***_*t*−1_), where the species population abundances ***n***_*t*_ at time *t* are given by some transformation *h*(***n***_*t*−1_) of the abundances on the previous time step. Specifically, it can be shown that the community matrix of such a complex process has eigenvalues identical to the MAR model matrix. The diagonal elements of ***B***, *b*_*ii*_, which represent the intra-specific, density-dependent effects, also satisfy the three existing theoretical definitions of the strength of density dependence^12^: the marginal effect on the per capita growth rate of an increase in density^28^, the derivative of the recruitment map at equilibrium ^28^ and the negative elasticity at equilibrium of the per capita population growth rate with respect to changes in the population ^29^. The latter measure is readily extendable to more complex life histories scenarios^29^.

Jointly, the model matrices ***B*** and **Σ** hold the key to formulating standardized measures of how a community reacts to environmental variation. These measures ultimately depend on the nature and intensity of the intra- and inter-specific interaction coefficients ^10,12,19,24^. Ives et al.^10^ derived four standardized metrics based on the ***B*** and **Σ** matrices and their eigenvalues. Variance Proportion (VP) quantifies how the long-run variance of the population compares to the variance of the environmental noise process. It is a summary of how the environmental noise distribution in blue in Figure 1 compares to the population size distribution in gray in Figure 1. As Figure 2 shows, differences in variability in the multi-species time series can be directly attributed to species interactions. In a stable system, the interactions among species that modulate changes in population sizes in a community from one generation to the next will be such that they cause the variance of the population abundance to be only slightly larger than the variance of the environmental noise (see Figure 1 for an example with a one-species system). On the other hand, in a less stable system, the species interactions greatly amplify the environmental variability, thereby generating large population fluctuations^26^. This amplification can be directly measured by the eigenvalues of the matrix ***B***, namely, by det(***B***)^(2/*p*)^ = (*λ λ* … *λ*)^2/*p*^ where *p* is the number of species in the system (see Ives et al.^10^ eq. 24 and subsequent paragraph). In the face of environmental variation, the growth rate of a population will react. This reaction is modulated, or filtered, by the intra-specific and inter-specific competition coefficients.

The Mean Return time (MR) and the Variance Return time (VR) refer to the amount of time it takes the system to return to its stationary distribution. It’s the stochastic equivalent of the deterministic return time. Specifically, it refers to the rate at which the transition distribution of the system converges to its stationary distribution. The shorter the time, the more stable the community is. Finally, Reactivity (R) is a measurement of how far the system pushes away from its equilibrium after it is perturbed and as Ives et al. argue, can be computed in two different ways, giving a total of four metrics of stochastic stability.

### From statistical ecology theory to practice: insights from a case study

In what follows, we applied the theoretical insights described above to an extensive data set of dynamic vaginal microbial communities. We then contrasted the resulting inference with the traditional practice of using the presence of a particular bacterial species at specific abundances from a snapshot of a bacterial community to imply etiology. We contend that such practice may, in the end, obscure rather than illuminate our understanding of the effects of different bacterial community compositions simply because population abundances can and often do vary widely over time. Additionally, we show how the concepts explained above contribute to answering questions of practical interest. For example, under which ecological scenarios (*i*.*e*., set of inter-specific and intra-specific interactions) will the abundances of species in a community quickly return from their current state to one where variation and composition regimes imply low health risks. How can the concept and measurement of “stochastic stability” contribute to estimating persistence probabilities?

The data we analyzed to exemplify the application of statistical ecology concepts were part of the Human Microbiome Project funded by the National Institutes of Health in which 135 women (see Clinical Study Methods in Supplementary Material). Women enrolled in this study self-collected daily mid-vaginal swabs for 10 weeks. We examined temporal changes in the composition of vaginal communities established using 16S rRNA gene sequencing. Every day after swab collection, each participant also measured vaginal pH (see Ravel et al. ^14^ for pH measurement methods). A simple examination of the temporal variation of pH in these samples (Figure 3) clearly illustrates that ample temporal variation in the dynamics of bacterial populations and their metabolic activities was the rule, rather than the exception. Important feedback loops between pH levels and bacterial metabolic activity are expected^14^ and these processes can be examined with our theoretical approach, as we explain below.

**Figure 3.**
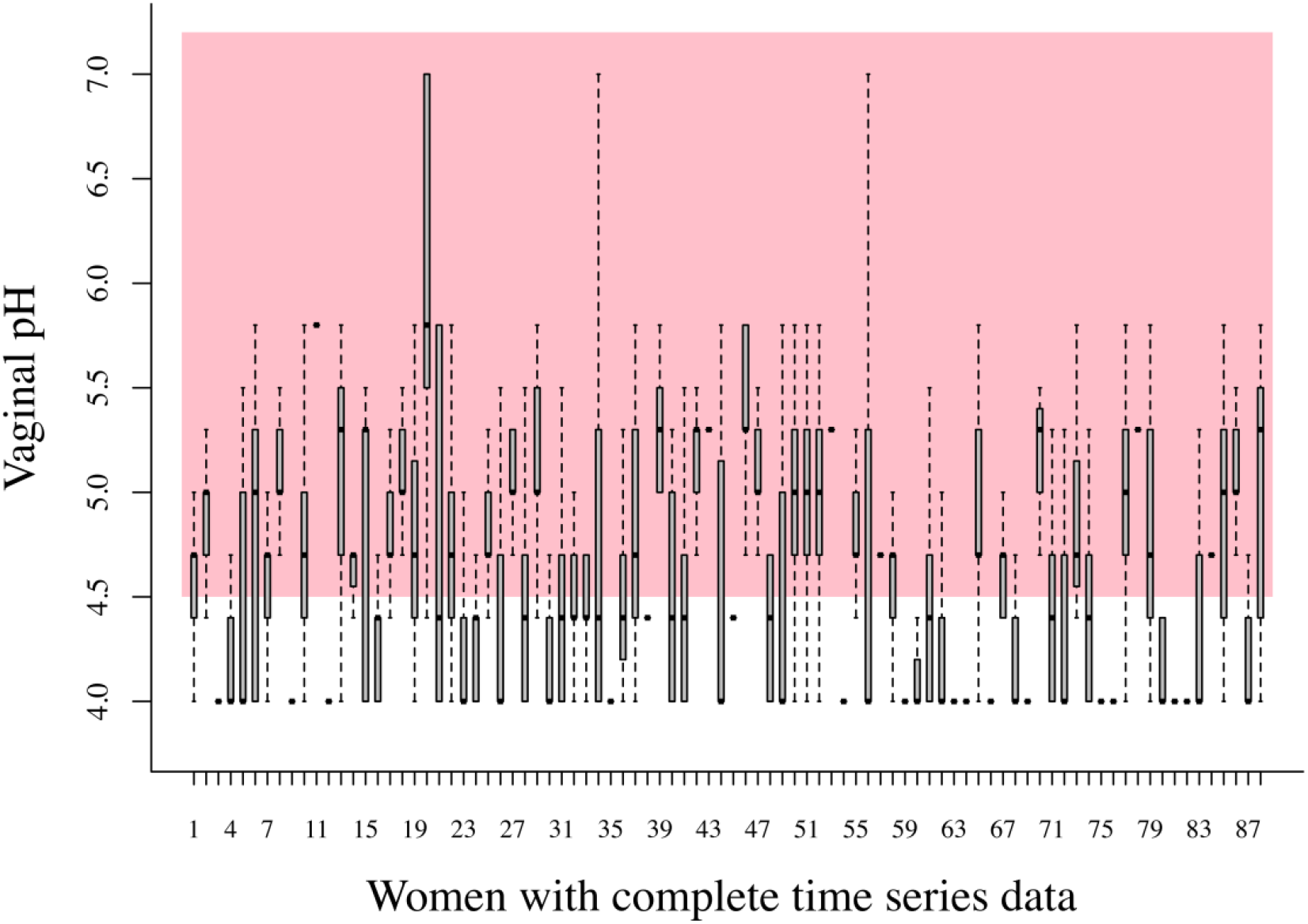
Boxplots of all the pH measurements taken over 70 days for 88 women in our bacterial community time series data, which were complete enough to estimate the MAR model parameters, as illustrated below. The boxplots here showcase the fact that the vaginal pH of 88 healthy, asymptomatic women varied widely over 70 days, inside and outside what is considered to be a healthy vaginal pH region (shaded pink and region < 4.5).

### Minimal sample sizes to fit a MAR model

Using the diagnostic tool presented in the Methods section, we determined which time series data sets had enough data to estimate the MAR model parameters. Computing the ratio of available data to the number of parameters to be estimated, we determined that 88 community time series out of the 135 total available could be reliably used for a full, multi-species population model-fitting analysis. The rest of the analyses are based on these 88 community time series data sets.

## Statistical decomposition of the sources of temporal variation

### Estimating the interaction coefficients: compositional data vs abundance data

We proceeded to fit the MAR model using the estimated population dynamics time trajectories without sampling error for all species and all data sets (Figure 4). While doing this second fit, the statistical uncertainty from the first step was propagated via parametric bootstrap.

**Figure 4.**
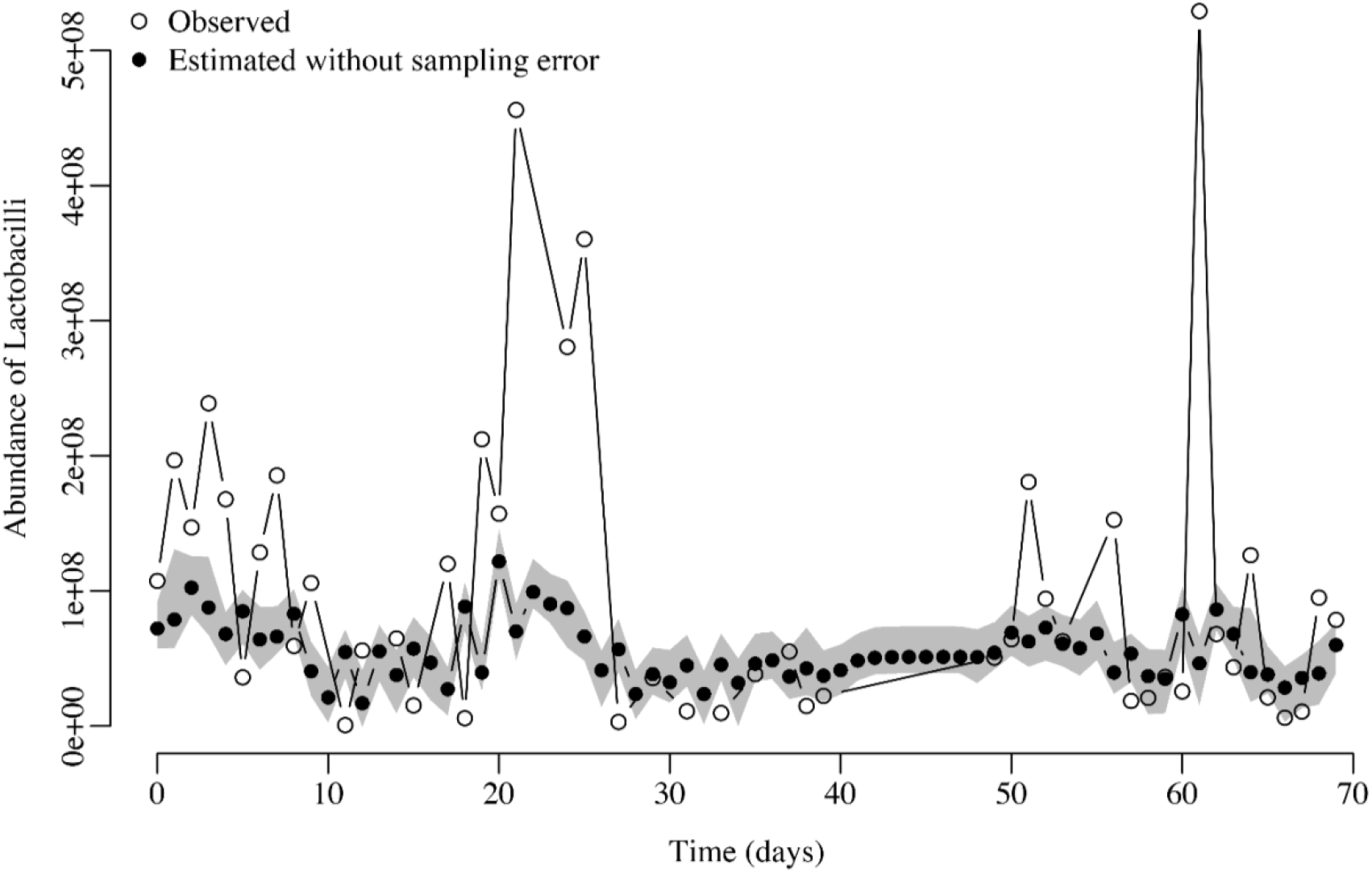
Observed *Lactobacillus* species abundances with error (empty circles) vs. estimated abundances (black circles) after accounting for sampling error for one time series. The gray area shows the 95% confidence interval of the estimated true abundances.

However, for this second step, microbiologists usually face a key practical decision: should they only work with relative abundances of species or work with both relative and estimated total abundances? For our case, the total abundance is the number of 16S rRNA gene copies per sample established using quantitative PCR. To make an informed decision as to which approach to undertake, here we simulated time series community abundances under the four scenarios shown in Figure 2. We then verified which interaction strength estimates were less biased (whether those resulting from using the compositional data or those resulting from using the total abundances). Although the relative abundances of species in a community (*i*.*e*., compositional time series data) are sometimes the only time series data available, our simulations showed that using compositional data leads to biased estimates of the interaction strengths specified in the matrix ***B*** of the MAR model (See Supplementary Information).

Our simulation approach was as follows: first, we selected the four community scenarios described in Figure 2 and simulated for each case 1000 time series of the abundance of the three taxa A, B and C. We then estimated the interaction coefficients using the MAR model described above fitted to both the relative abundance time series and the absolute abundance time series. The absolute abundances were estimated by anchoring the proportions into total abundances at each time step. Next, we fitted the MAR model to estimate the interaction coefficients using the 1000 time series of relative abundances and the 1000 time series of total abundances. Then, we calculated the ratio between the estimated and the true interaction strength in each case. When the interaction coefficients were estimated appropriately, a boxplot centered at 1 with a small variance resulted. We estimated if each ratio between estimated and true coefficients departed from an expected value of 1. The results of this simulation experiment (see Supplementary Figures 1-4) clearly show that when the total abundances are used, the relative bias boxplots are centered around one. When compositional data is used, those boxplots have a much wider interquartile range and most of the time, are not evenly centered around one. Thus, fitting the MAR model to compositional data tends to lead to severely biased estimates of the interaction strengths. Therefore, the best approach to estimate the interaction coefficients is to use total abundance data.

In longitudinal studies of microbiomes, the number of 16S rRNA gene copies only provides estimates of the absolute abundance of taxa and not the true abundance of each bacterial species. Our simulations demonstrated that estimating the strength of intra-specific and inter-specific interactions based on relative abundance data results in biased estimates of the interaction strengths. Hence, we performed a pan-bacterial qPCR assay to quantify the total 16S rRNA gene copies in each sample, which estimates the absolute bacterial abundance in each sample. Estimates of true abundance were then calculated for each taxon by multiplying relative abundance by total 16S rRNA gene copies. The qPCR assays were done in triplicate for each of the 135 women to document the variability in species abundances due to observation error.

We fitted the three different multi-species population dynamics assemblies/model variants using the MAR model of Ives et al. ^10^ and the de-noised time series data sets. The first model variant used all 13 species mentioned above. The second model variant required fitting a three-species model where we grouped all four *Lactobacillus* species into a single ecological species, *Gardnerella vaginalis* as the second species, and the other eight species grouped into a third species. For the third variant, we fitted a simple 2-species model with all four *Lactobacillus* species grouped as the first species and the other nine species grouped as the second taxon.

We computed Ives et al. stochastic stability metrics with the MAR model parameter estimates for each model variant (13 species, 3 species, and 2 species models). For each one of the three cases, we then classified the 88 vaginal bacterial communities into four different stability categories using a Principal Components Analysis (PCA) on their estimated stability metrics. Using k-means clustering on the resulting PCA scores for these women and the fact that for all these metrics, lower values indicate higher stability, the 88 bacterial communities were classified into four different categories: Highly stable, stable, unstable, and highly unstable. The best classification scheme out of the three different multi-species models corresponded to the two-species MAR model, where we pooled all 4 *Lactobacillus* species into the first taxa and the other 9 species into the second species (the performance criterion to pick the best classification scheme was the amount of variance explained by the analysis). This classification scheme is shown in Figure 5.

**Figure 5.**
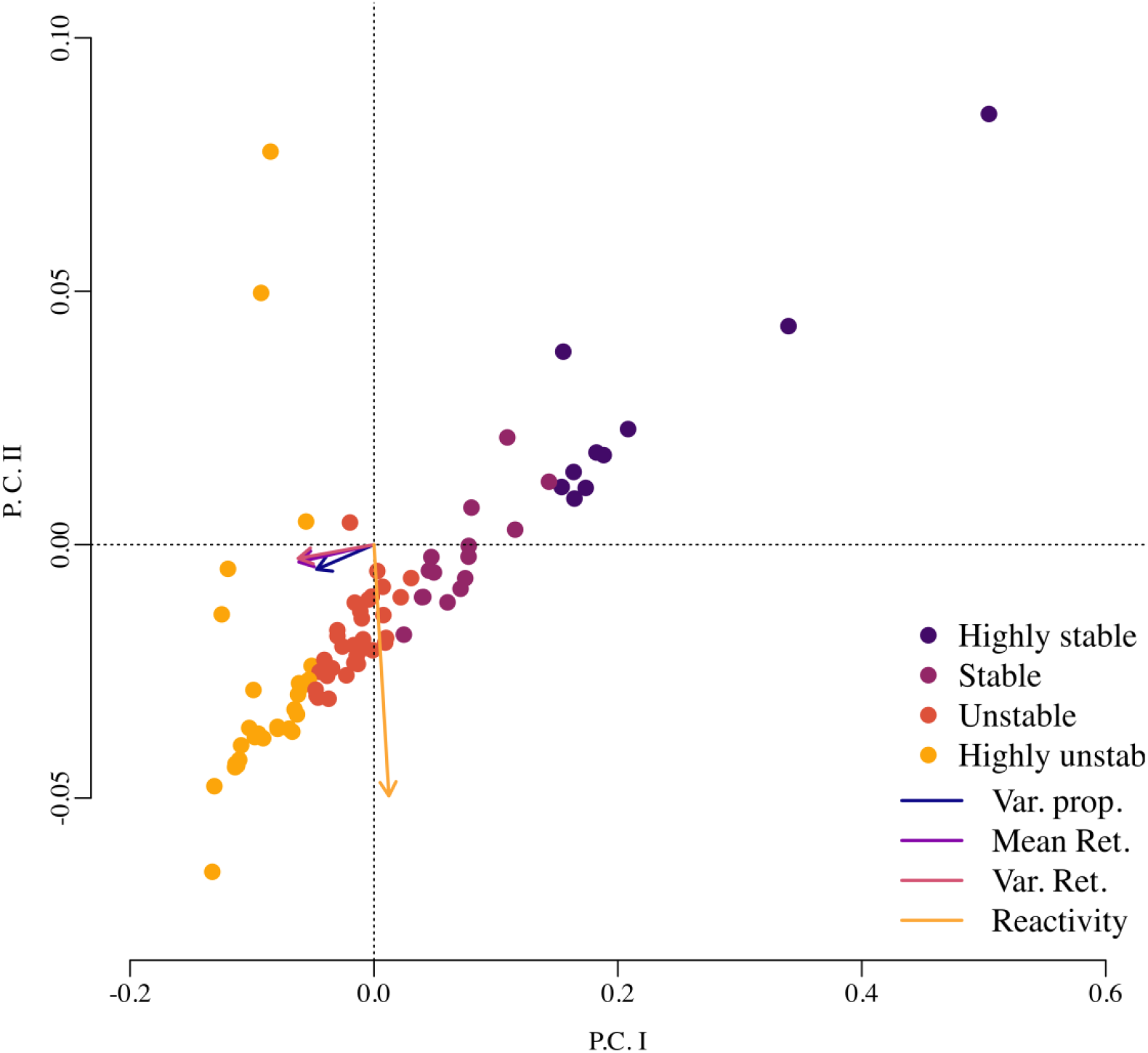
Principal component analysis (PCA) was performed on the four stochastic stability metrics estimated for the vaginal bacterial community time series data of 88 women. In this analysis, the samples (rows) correspond to each woman, and the four columns (variables in the PCA analysis) correspond to the four stability metrics estimated by fitting the MAR model of Ives et al. The arrows’ lengths and direction represent the strength of association of each one of these four metrics with the principal component axes: the variance proportion, the mean return time, and the variance return time are highly associated with the first principal component, while the reactivity is highly associated with the second principal component. Using k-means clustering, the PCA scores of these 88 bacterial communities were classified into four groups. Because lower values in these stability metrics indicate higher stochastic stability, an examination of the magnitude of these four metrics in each one of these four groups suggested the labeling of highly stable, stable, unstable, and highly unstable dynamics (see Supplementary material for details).

### Estimation of persistence dynamics from the MAR model parameter MLEs

Alone, our stability classification scheme is an insufficient approach to understanding multi-species population dynamics because the “stable” and “unstable” attributes are given here to a community without regard for the health risks associated with its composition. Stability, a property of dynamic systems, should not be equated with desirable or undesirable behavior in terms of health outcomes because one community can have stable population dynamics but sustain a low relative abundance of a strain, thus bringing high health risk. Thus, considering overall abundance and composition in addition to stability is needed to assess the desirability of a particular community dynamics. Indeed, Klatt et al ^30^ show that when the relative abundance of *Lactobacillus* dwindles below a 0.5 proportion, the bacterial community is under a high risk of infection by HIV. On the other hand, as the relative abundance of *Lactobacillus* moves above 0.5, the risk of infection decreases. Seeking to elucidate which type and magnitude of ecological interactions would lead to desirable dynamics (i.e., fluctuations in the relative abundance of *Lactobacillus* above 0.5) is a reachable target under our analysis using the MAR model. If attaining a sustained high relative abundance of *Lactobacillus* over time is a health-management target, as in Klatt et al ^30^ (see Supplementary Figure 5), then we contend that our approach described next should be used.

We developed a Risk Prediction Monitoring (RPM) tool that estimates the temporal changes in persistence probabilities. This method mirrors conservation biology approaches for population monitoring in which a metric measuring extinction risks is periodically updated with any change in the population called Population Viability Monitoring, or VPM ^31^. Before explaining and implementing our RPM tool, we first explain how the well-known VPM method from Staples et al. works and apply it directly to one of our 88 data sets to exemplify it. Immediately afterward, we fully develop and implement our RPM method.

The VPM method consists of serially estimating the persistence probabilities with every data point added to the current length of the time series of population abundances. For annually reproducing species, with every year that passes, a new total abundance is recorded. With it, an updated estimate of extinction risk is computed. Repeating the same process for multiple years yields a temporal trend of extinction risks. Consider the following example from conservation biology: if population abundances of a threatened species have been available for the past 30 years, and if managers want to check whether, as of late (*e*.*g*., for the past 10 years), the extinction risk of the population has been increasing or decreasing, then the following is done ^31^: First, a stochastic population dynamics model is fit using the first 20 years of the data. With the model parameter estimates and the data up to year 20, the probability that the population will crash below a critical threshold within the near future, *e*.*g*., during the next 5 years, is computed. The resulting probability is recorded. Next, the observed population size for year 21 is added to the time series. The model parameters are then re-estimated, and the probability that the population will crash sometime during the next 5 years after year 21 is computed. That probability is also recorded. Iterating this process for 10 more years yields a time series of extinction risks for the last 30 years.

Here, we exemplify the conservation biology method with one of our 88 bacterial community datasets. For our vaginal bacteria data set, our target was to track the probability that the proportion of all the *Lactobacillus* species in a vaginal bacterial community drop below 50%. Our time unit, in this case, is days, as new swabs were collected daily. The abundances for all bacterial taxa and the proportion of *Lactobacillus* species were available for 70 time-steps. In Figure 6, upper left panel, we first fitted the stochastic multi-species Gompertz model of Ives et al ^10^ to a single time series of observed abundances and proportions of *Lactobacillus* up to day 30 (black empty circles). We did so by placing all *Lactobacillus* taxa as one type in the model, and all other species were pooled together as a second type. We then used the model parameters to project in the next ten days the *Lactobacillus* abundances and their proportion in the population 50,000 times (grey lines). The proportion of such projected trajectories that dropped below 50%, which was 0.4 in the upper left panel, is an estimate of the *Lactobacillus* persistence probability above 50% during those ten days. With every passing day, this estimate was updated. In the next three panels (upper right, lower left, and then lower right), we show these simulations for only days 40, 50, and 60, but daily changes in persistence probabilities for days 30 to 70 were computed.

**Figure 6.**
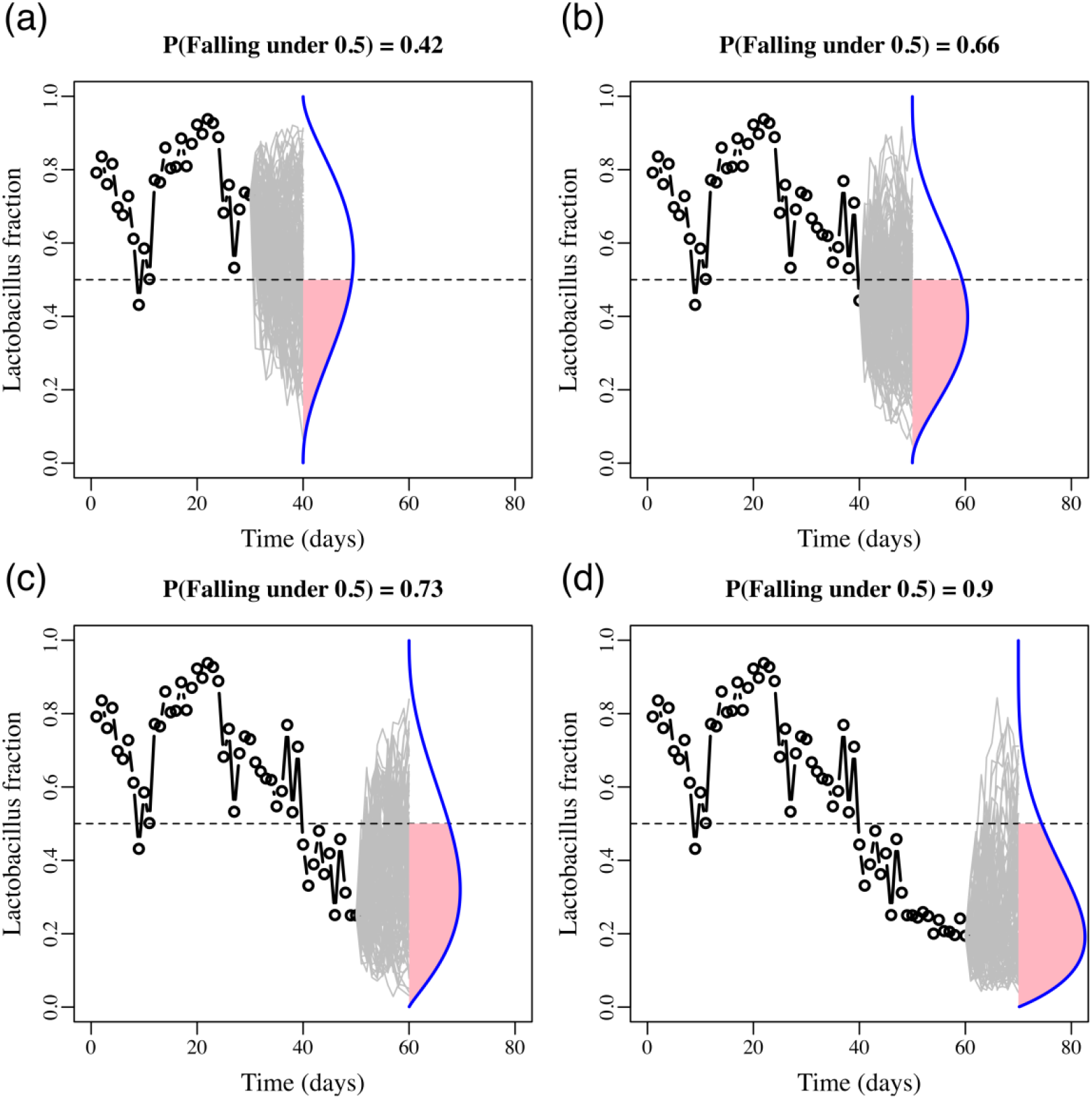
Population viability monitoring and estimating the temporally varying chances of *Lactobacillus* persistence. Illustrated is an example of Risk Prediction Monitoring (RPM) using stochastic population dynamics models.

The VPM method illustrated above for our data set is essentially retrospective, but here we devised a prospective modified version of it that allows for the comparison of the dynamics of multiple communities in the near future. Furthermore, we switched the estimation focus from tracking the probabilities of crashing below a population size or proportion threshold to following their complement, persistence probabilities. This modified method links our stability metrics with the risk assessment task and is called the RPM tool. We developed the RPM tool because we faced the problem of assessing the risk dynamics for all 88 communities and being able to evaluate these under the same level playing field. To make such comparisons, we evaluated the risk dynamics while answering the question: How would the risk of *Lactobacillus* spp be falling below 45% change over the next 20 days if all communities were started with the same proportion (50%) and then monitored over the next 20 days? We answered this question by implementing these steps: First, we retrieved the MAR model parameter estimates for all 88 communities. Using these estimates, we computed the MAR model predicted mean abundance of *Lactobacillus* at stationarity for every case. We then set these mean abundances as the starting abundances for a 20-day projection in each case. Additionally, we assumed that the starting total abundances for the non-*Lactobacillus* taxa in all these projections were equal to these abundances. Thus, if, in one case, the mean abundance at stationarity of *Lactobacillus* was predicted to be 3.5x10^8^ 16S rRNA gene copies per swab, the starting mean abundance of the non-*Lactobacillus* species were assumed to be identical, 3.5x10^8^ 16S rRNA gene copies per swab. With these starting values, we computed the mean projected abundances for the next 20-day trajectories. We used these to numerically estimate via simulations the probability that the *Lactobacillus* taxa would remain above 45% on day *t* for *t* = 1,2,3, …,20. The resulting trends in *Lactobacillus* persistence probabilities are shown in Figure 7.

**Figure 7.**
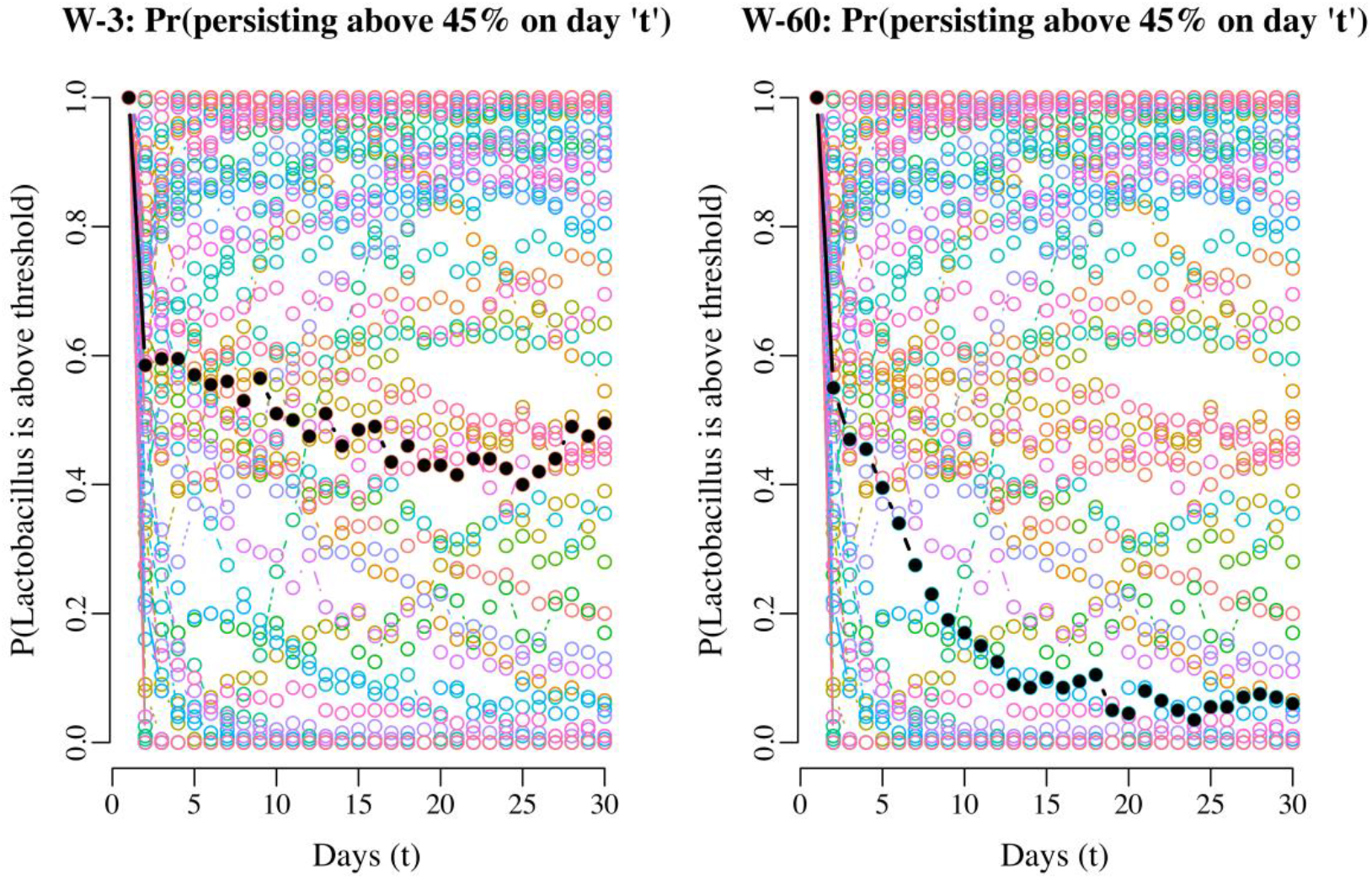
Projecting the probability of *Lactobacillus* persisting above 45% for two women, starting at 50/50 from their carrying capacities for 20 days. The black dots are the projected probabilities of persisting for woman 3 (on the left) and woman 60 (on the right), for 30 days. As a background and in different colors, the same trend is shown for all the other women in the study. The wide array of trajectories of all the trends for the other women emphasizes the wide variability in predicted community dynamics.

Our RPM tool was used in conjunction with the estimated matrix ***B*** to identify which interaction coefficient drove each persistence trend. We found that a decaying persistence trend of a bacterial type of interest was explained by whether other bacteria impacted its growth negatively or positively, which is the information contained in ***B***. We illustrated this finding using the resulting decaying RPM trend for woman 60 (right panel of Figure 7). The estimated two-by-two matrix of interactions ***B*** for woman 60 is as follows: the one-step total effect of non-*Lactobacillus* species on the per capita growth rate of *Lactobacillus* species had a negative coefficient, -0.39. On the other hand, *Lactobacillus* species had a small positive effect on non-*Lactobacillus*, 0.001. Therefore, the presence of non-*Lactobacillus* taxa had a negative density dependence effect on the growth of *Lactobacillus*, while *Lactobacillus* had a positive effect on the growth rate of non-*Lactobacillus*. In the end, this asymmetry negatively affected the growth of *Lactobacillus*. In both cases, the strength of the intra-specific density dependence was weak (0.86 for *Lactobacillus* and 0.76 for non-*Lactobacillus* species). To verify whether that asymmetry in the inter-specific growth rate effects was what drove the decay in persistence probabilities for the *Lactobacillus* taxa, we did two numerical experiments: for the first experiment, we switched the sign of the effect of one group on the other, so that non-*Lactobacillus* had a positive effect of 0.39 in the growth rate of *Lactobacillus* and in turn, *Lactobacillus* had a negative effect of -0.001 on the growth rate of non-*Lactobacillus* taxa. Next, we re-computed the RPM trend in persistence probabilities using this modified ***B*** matrix of interactions. The resulting RPM trend of persistence probabilities, plotted in pink in Figure 8, remained at 1 for the next 20 days. The second numerical experiment consisted of artificially increasing the maximum growth rate of the *Lactobacillus* taxa while leaving the ***B*** matrix of interactions unchanged. Then, a restored trend in persistence probabilities was also obtained (Figure 8).

**Figure 8.**
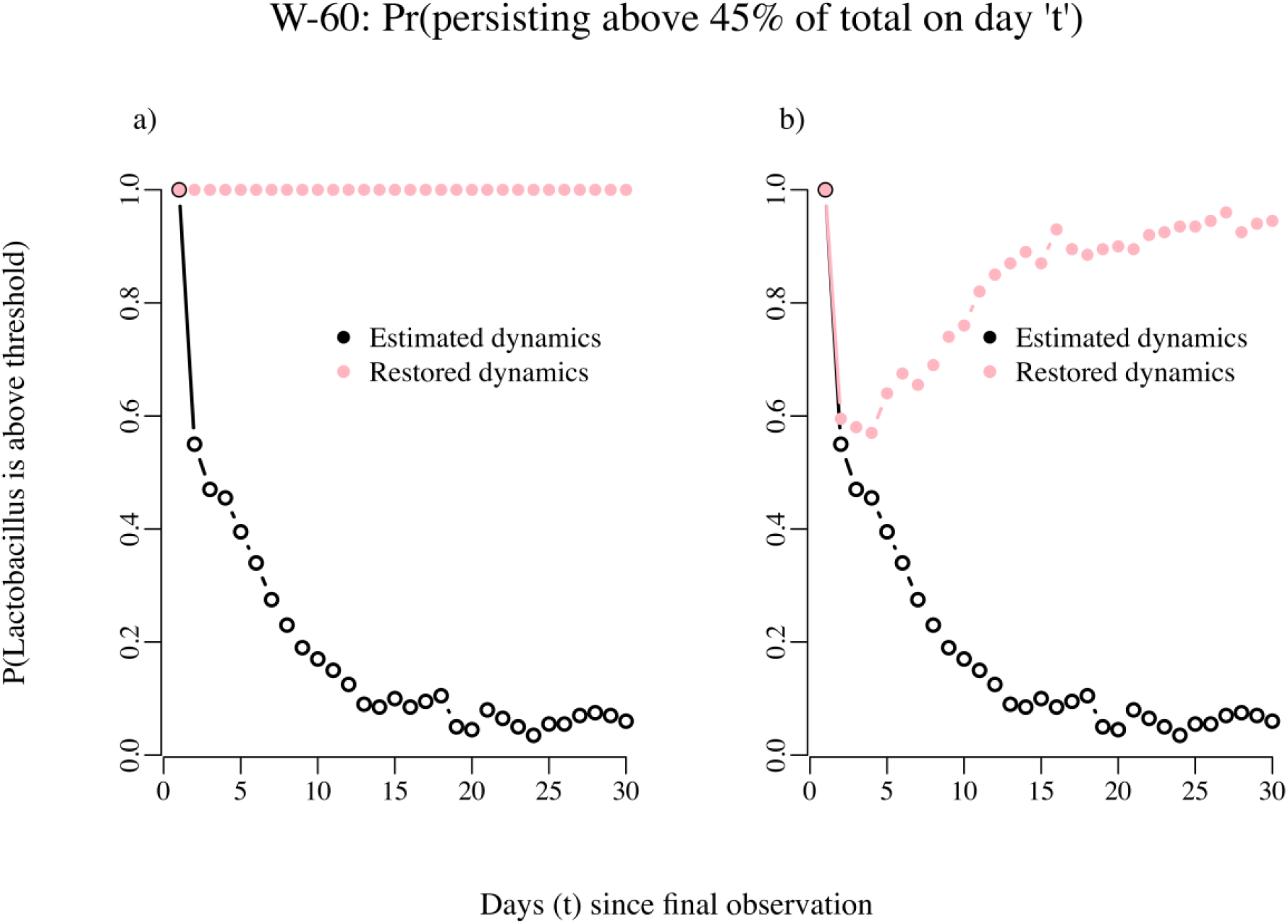
Probability of persisting above 45% of total abundances “t” days into the future. As explained in the text, the changes in the estimated coefficients of the interactions matrix results in restored dynamics and persistence probabilities. In panel A the effect of *Lactobacillus* on non*-Lactobacillus* was changed from positive to negative and the effect of non*-Lactobacillus* on *Lactobacillus* was switched from negative to positive. In panel B only the maximum growth rate of the *Lactobacillus* species was increased by 25%.

## Discussion

Our case study revealed clear and quantifiable differences in the stability and persistence dynamics of individual vaginal microbial communities. These results extend beyond abstract ecological principles and offer biologically meaningful insights into the behavior of bacterial taxa that are directly relevant to women’s reproductive health. First, we classified vaginal microbiota using inferred ecological interaction strengths into four distinct stability categories: highly stable, stable, unstable, and highly unstable. Importantly, these categories were not based on species composition alone but on ecological processes governing community dynamics, such as density dependence and interspecific interactions. This reframing allows for a deeper understanding of what it means for a vaginal microbiota to be stable, moving beyond simplistic assumptions equating *Lactobacillus* dominance with health. Second, by linking the MAR model outputs to a practical tool, our Risk Prediction Monitoring (RPM) framework, we provided individualized, time-resolved estimates of the persistence probabilities of *Lactobacillus* spp. This allowed us to identify vaginal microbiomes at risk of transition to low-*Lactobacillus* states and to pinpoint the ecological interactions (*e*.*g*., asymmetric competition from anaerobes) that drive these transitions. Third, we showed through simulations that specific changes in ecological parameters, such as reducing the negative impact of *non-Lactobacillus* species or boosting *Lactobacillus* intrinsic growth rates, can theoretically restore persistence probabilities. These insights are directly translatable to microbiome modulation strategies aimed at promoting resilience and favorable community dynamics, whether through probiotics, prebiotics, or ecological engineering. Our modeling and case study demonstrate that the ecological architecture of vaginal microbiota, and not just their taxonomic composition, dictates their stability and susceptibility to shifts associated with increased disease risk. By explicitly connecting microbial ecology to clinical relevance, we provide a rigorous and accessible framework for understanding and managing the vaginal microbiome.

This study reflects an unprecedented integration of ecological, mathematical, statistical, and conservation biology in which principles to understand and predict the dynamics of an extensive microbial community’s time series data set. In the past few years, the complex nature of microbiome data has brought together an ever-growing number of multi-disciplinary research teams ^32^. Yet, the fast pace of modern methodological research in microbiome studies contrasts sharply with the paucity of population dynamics studies seeking to understand from basic principles the benefits or shortcomings of novel data analysis techniques. The main motivation of this study was that, by and large, variability in microbial time-series data is still perceived as “statistical noise” rather than as an intrinsic property of the growth of bacterial communities. Phrasing through the MAR model variability over time as an intrinsic property of a growing population allows linking concepts like the strength of intra-specific and inter-specific competition to the qualitative response of a population in the face of uncertain environments. Not only can these competition coefficients be estimated, and the stability of the system can be assessed by fitting the MAR model, but the chance of persistence of bacteria taxa can also be further assessed. To our knowledge, this is the first study demonstrating how persistence probabilities of medical and ecological interest bacteria can be estimated and even manipulated by identifying which interaction coefficient strengths are their main drivers. We thus demonstrate how the apparently simple stochastic multi-species time series model of Ives et al. (2003) can be used beyond its original applications to approach some of the most pressing questions regarding monitoring bacterial communities ^8^.

Vaginal communities dominated by species of *Lactobacillus* have been associated with health and a reduced risk to diseases such as bacterial vaginosis or sexually transmitted infections. The notion that the dominance of *Lactobacillus* is associated with health is deeply ingrained in the field of women’s urogenital health and strongly supported by the findings of numerous studies ^33,34^. Regrettably, the converse — that low proportions or the absence of *Lactobacillus* is unhealthy — has also permeated the field’s lexicon. This is a logical fallacy of denying the antecedent ^35^, which essentially argues that if healthy women have vaginal communities dominated by *Lactobacillus*, then the absence of *Lactobacillus* in vaginal communities is, in itself, unhealthy. This claim is refuted by the findings of numerous studies on the species composition of healthy, asymptomatic women that have shown that a significant proportion of healthy asymptomatic women have vaginal communities with low proportions of *Lactobacillus* ^36–38^. Instead, they are dominated by various species of strictly and facultatively anaerobic bacteria, such as *Gardnerella vaginalis, Mobiluncus, Prevotella, Brevibacterium, Peptoniphilus*, and others (Onderdonk et al 2016). With that said, it should also be recognized that low proportions of *Lactobacillus* in vaginal communities are associated with an increased risk of disease. However, it is not a disease state *per se*. Nonetheless, investigators have often referred to these communities as abnormal ^39^, out of balance ^40^, or in a state of dysbiosis ^41^ that needs to be corrected. We posit that except for symptomatic bacterial infections, all other states are ‘healthy’, and in many instances, they are ‘normal’ (meaning they are often observed). However, they may differ in terms of disease risk, and modulation should be considered in that context.

Most studies on the species composition of vaginal bacterial communities have employed cross-sectional designs that yield point estimates of community composition. It seems to be assumed that the species composition of communities is relatively invariant over time in the absence of natural or unnatural environmental disturbances such as menstruation or the use of lubricants ^42–44^. Contrary to this assumption, longitudinal studies have shown that the vaginal microbiota of many women is dynamic and often transitions through states in which *Lactobacillus* spp. are lacking ^42,45^. These states vary in frequency and duration and are therefore associated with varying levels of risk for urogenital infections and other maladies. One could reasonably consider these to be windows of elevated risk that can open and close, sometimes over very short periods.

Our PCA and MAR model-based stability classification scheme (Figure 5) takes a first, admittedly imperfect step toward process-based management of bacterial community dynamics and rigorous use of the term “stability”. Although previous community classification schemes using PCA relied on patterns of abundances, our approach relies on inferred ecological processes from the time series of abundances. As theory and current practice in conservation biology show, the longer the multi-species time series data, the better the information regarding species interactions in a community can be better teased apart. We went one step further and estimated how these inferred interactions ultimately govern the community response to environmental variability. The nature of such response was quantified with Ives et al.’s (2003) four stability metrics, and the PCA in Figure 5 separates bacterial communities according to these metrics (see Supplementary material). Thus, the position of each bacterial community in the PCA space is determined by the strength of ecological interactions. If one community is largely unstable, an analyst can peer into the nature and intensity of those estimated interactions and change them one by one to move the community in PCA space from an unstable group into another classification group. In other words, an investigator can test statistical hypotheses regarding which interactions are responsible for one or another stability classification result. Identifying interactions that render a community stochastically stable can be the first step in a research agenda that seeks to understand how to guarantee such stability by modulating the strengths of interactions. Our RPM approach is a natural extension of our stochastic stability inferences. It is an easy-to-understand approach to approximate the time-dependent persistence probabilities of the bacterial species of interest. As Olesen and Alm ^40^ have argued, tools like our RPM approach that focus on prediction rather than simply detecting differences are needed, and here we deliver on that particular need.

Using this persistence probability methodology in studies of the vaginal microbiome would mirror an approach called Population Viability Analyses that has been successfully used in conservation biology for many years ^12,46^. Unlike the majority of cases in conservation biology our model choice (Ives et al’s MAR model) has been extensively tested in a recent theoretical-simulation study^12^. Estimates of the strengths of interactions can be used to formulate hypotheses regarding the molecular mechanisms and genetic composition that underpin different types of interactions. By plotting the variability in the sign (positive or negative) and intensity of interaction coefficients (for example, the effect of *Lactobacillus* on the growth rate of *Gardnerella* or some other species) one can locate and isolate cases where the sign of species interaction relations flip (say from positive to negative) and eventually guide the laboratory determination of the genetic composition of strains associated with interaction relationships in every quadrant (Supplementary Figure 6).

Adopting statistical ecology theory and concepts reveals the inconsistencies of using terms like “dysbiosis” to characterize a microbial community. Dysbiosis is commonly defined as a change in the composition and function of a human microbial community that is typically driven by environmental and host-related factors that exceed a community’s resistance and resilience ^47–49^. However, this definition doesn’t fit fully with what theoreticians in ecology understand as resilience and resistance. Resilience, on the one hand, is the rate at which a community returns to a state that existed prior to a change. Conversely, resistance is the magnitude of a community’s response to a given disturbance ^50^. Both resilience and resistance are built into Ives et al stability metrics. Instead of trying to frame a dysbiosis definition into these concepts, it seems much more straightforward to use Ives’ stability metrics directly to classify the stability dynamics of a community, just as we do here. Additionally, in current practice, investigators often state that ‘healthy’ communities are ‘in balance’ ^40,51,52^. This terminology reflects an erroneous assumption that the composition of bacterial communities in healthy individuals is essentially invariant and that changes in the relative abundances of species are necessarily bad and, in some cases, constitute sufficient evidence to classify these variants as disease states. This classification is often done based on pairwise microbiome comparisons at two points in time. Except for symptomatic bacterial infections, all other states seem ‘healthy’, and in many instances, they are ‘normal’ (meaning they are often observed). These words and phrases are loosely defined and inconsistently used, and leading to confusion among non-experts. The literature is peppered with examples ^40,51,52^. Instead of seeking a definition of dysbiosis, we assert that it might be better to translate theoretical ecology concepts into practical approaches for the management of human-associated bacterial communities. This can be accomplished using concepts and methods that have come to be well-known in the fields of population dynamics and conservation biology.

Population dynamics as a field in ecology has long touted the theoretical and practical advantages of jointly modeling demography, and the influence of the environment and sampling error ^15,19^. At the same time, conservation biology has taken advantage of these ideas and modeling approaches to predict population persistence probabilities ^31,46^. Here, we have shown that the same sort of stochastic population dynamics equations can be used to re-phrase the concept of stability as the magnitude of the reaction to a variable environment. Our work represents the first comprehensive integration of theoretical stochastic population dynamics, unusually long time series of bacterial community abundances, and conservation biology principles. This integrated approach resulted in two major steps towards a better understanding of human-associated bacterial communities. First, by estimating each bacterial community’s reaction to exogenous variability we achieved a stability-based ecological community classification. Second, we provide estimates of the short-term persistence probabilities of bacterial types of medical interest for the first time. This result is important because our estimated temporal trend in persistence probabilities can be used to construct an evidence-based inference regarding the fate of a pathogen, for example. Finally, we conclude that a comprehensive examination of the reach of stochastic population dynamics modeling in microbial community ecology is beginning to take shape as a body of work. Our efforts provide a theoretical framework that can very well represent microbial phenomena of interest in a simpler and unified way as effects of a common cause: an alteration of the growth rate of a population by itself, by another population or by the environment.

## Methods

Fitting the MAR model to extensive time series of microbial abundances presents at least three major methodological challenges: The first is determining whether there exists enough information in the data to estimate the MAR model parameters. This question boils down to determining which time series length is sufficient to provide statistically sound parameter estimates. The second methodological task is separating the environmental process variability from sampling noise. The third one deals with time series data that are incomplete in these ecological studies. In what follows, we detail our approach to these three problems. We note that all human subject research studies were performed at University of Maryland under approved protocols by the Institutional Review Board of the University of Maryland Baltimore under numbers HP-00040935 and HP-00041351. Samples were collected after obtaining written informed consent from all the participants, and all experimental methods complied with the Helsinki Declaration.

### Minimal sample size to fit a MAR model

The quality of the statistical fit of the MAR model depends on the amount of information present in the multi-species data set. This information can be measured through the statistical properties and diagnostics related to the model parameter estimates. The statistical quality of the parameter estimates aresin turn related to how many data points per parameter, or “degrees of freedom” one has available to do model fitting. Another way to think about quantifying this information is by computing the ratio of data to the number of unknown parameters. Ives et al model is, however, quite data-hungry: Let *p* be the number of species in the data set. The vector of maximum growth rates *A* has *p* unknown parameters. The matrix of interactions *B* and the variance-covariance matrix of the environmental fluctuations Σ have each *p* × *p* unknown parameters, thus, the total number of model unknowns is 2*p*^2^ + *p*. With 13 species this number is 351. This number can be compared to the available number of independent data points in order to gauge if one has enough “degrees of freedom” for estimation.

Because this model is Markovian, every time-step transition (change in population abundance) is an independent data point. The likelihood function of the MAR model, from where its parameter estimates are derived, is therefore computed as the product of all the observed transitions. This likelihood is maximized to obtain the parameter estimates. If *n* is the length of the time series (70 in our case, see below), then the number of transitions that can be used for the maximization of the likelihood function is *n* − 1. If *m* is the number of replicated samples per species per time point, then the number of data points available for parameter estimation is simply (*n* − 1)*mp*. Consequently, for the estimation to be feasible, one needs to verify that (*n* − 1)*mp* > 2*p*^2^ + *p*. On the other hand, solving for *n* in this inequality gives the minimum sample size (time series length and/or number of replicates per time step) needed to ensure estimability as 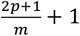, which is equal to 2(*p* + 1) in the common case where *m* = 1. For example, with 13 species and one replicated time series with no observation error, 2(*p* + 1) = 28 and the ratio of observations to number of parameters is 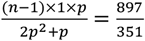.Finally, this thinking can be extended by including the parameters needed to decompose biological (process) variation from sampling error variation.

### Statistical decomposition of the sources of temporal variation

In this study, we decompose the changes in abundances of species over time into three of the four main components mentioned above ^4,19,23^: 1) Population growth, density dependence and inter-specific interactions, or predictable changes in births and death due to current and past abundances of the species in the system 2) environmental stochasticity or (random) temporal variation in vital rates representing variation in environmental conditions (good/bad times for survival and reproduction and 3) observation error and/or sampling noise which if left unaccounted can lead to grossly mis-represented dynamics ^19,23^. Demographic stochasticity, the fourth component, although not included in this first phase of our studies can be accommodated in time series estimation methods ^11^.

State-space models, widely known as statistical hierarchical models, allow decomposing the biological and sampling sources of variation using a one pass statistical fit ^19^. Stochastic population models with added observation error are just one example of this wide class of models. Although these models are routinely used ^13^, it has long been known that their fitting isn’t without statistical difficulties due to parameter identifiability problems, among others ^19,53^. Recently, statistical ecologists have extensively documented and demonstrated such challenges ^13^. Further studying the statistical and scientific merits of different computer intensive approaches to obtain either the maximum likelihood estimates (via Data Cloning, the Laplace approximation, the Geyer-Thompson likelihood ratio algorithm, Monte Carlo integration to name a few) or the Bayesian posteriors as well as Bayes Factors for these state-space models is a task that merits its own, separate efforts^54^ and goes well beyond the conceptual scope of this manuscript.

Our present approach to fit a multi-species population model to a bacterial community time series data set was as follows: first, we estimated the most likely location of the true, unobserved abundances with sampling error removed, along with their confidence intervals using the Kalman estimation methodology ^23^. This methodology simultaneously accounts for sampling error and missing data points in the time series of abundances. The resulting observation-noise filtered time series of abundances were then used to fit the MAR stochastic population dynamics model for the entire community. While doing this second fit, the statistical uncertainty resulting from the first observation error step was propagated via parametric bootstrap ^55^. Separating the estimation of the observation error from the biological process error allowed us to be sure at each step that the Mean Squared Error (MSE) of the model parameters were adequate via extensive simulations (github.com/jmponciano).

Parameters for the MAR model can be estimated with three methods: Maximum Likelihood (ML), Conditional Least Squares (CLS) or a Bayesian framework. The first two approaches were described for the first time for univariate (*i*.*e*. one species) in the landmark publication in Ecology by Dennis and Taper ^56^. Because the MAR model and its univariate version, the log-Gompertz model^19^ are written as a linear recursion, Dennis and Taper ^56^ and Ives et al^10^ showed that CLS estimation amounts to obtaining the value of the model parameters that minimize the squared difference between the observed population abundances at each time step and those predicted by the model, conditional on the population abundances at the previous time step. Thus, simply regressing the log-abundances at all times *t* with those at times *t* − 1 results in a very fast, MCMC free estimation of the Gompertz population dynamics model with environmental noise. Whereas this minimization is achieved through a standard matrix algebra least-squares calculation for this regression, inference via confidence intervals and uncertainty propagation must be achieved via bootstrapping or another computer intensive method, like data cloning^53^, to account for the time dependencies between observations. Indeed, unlike a simple regression where observations are independent, here the log-abundances result from time dependencies stipulated by the population dynamics model.

Translated to a setting with *p* species as in the MAR model, the one-step ahead regression above leading to CLS estimates is written as a regression for each species of its log abundances at time *t* with its own species log abundances and those of all the other *p* − 1 species at time *t* − 1 (see^10^). Thus, for each species, a least squares calculation is required and hence estimation of the MAR model parameters is a simple iterative matrix algebra least squares calculation. The required matrix algebra setting for the multi-species MAR is as follows: starting with a community time series data with *q* + 1 time points and *p* species, let the matrix ***X*** of dimensions *q* × *p* contain in each row *i* the vector of log-abundances (after noise-filtering) for each one of the *p* species in the community, for time steps *i* = 0,1, …, *q* − 1. As well, let the matrix ***Y*** of dimensions *q* × *p* contain in each row *i* the vector of log-abundances (after noise-filtering) for each one of the *p* species in the community, for time steps *i* = 1,2, …, *q*. Finally, let ***E*** be the *q* × *p* matrix of process error terms. Then, the MAR model evaluated at the observations can be written as

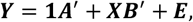

where **1** is a *q* × 1 vector of ones and ***A*** and ***B*** are respectively the vector of maximum growth rates for each species and the *p* × *p* matrix of interactions as defined above. Let ***Z*** = **[1, *X*]** be a *q* × (1 + *p*) matrix with the vector **1** and the matrix ***X*** concatenated horizontally. Then, setting as ***D***_*i*_ = [*a*_*i*_, ***B***_*i*_] as a 1 × (1 + *p*) vector with the maximum growth rate and the inter and intra specific coefficients for species *i* (**B**_i_ corresponding to the *i* ^th^ row of the matrix ***B*** containing the coefficients of the effect of every species on the per capita growth rate of species *i*) for every species the CLS estimates of the MAR model parameters 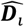 are given by 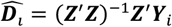, where ***Y***_*i*_ is the *i* ^th^ row of the matrix ***Y***. Finally, note that in our Github repository https://github.com/jmponciano/StochasticMicrobiome/tree/main/ExampleCalcs/R we give a fully commented R function for the MAR CLS calculation and for all the stochastic stability calculations. The function for the MAR CLS implementation is *mar*.*cls()* and the function to compute the stochastic stability calculations is *stability()*. These correspond to the translation to the language R of the original functions coded by Ives et al. Finally, note that in our repository as well we provide three fully commented html vignettes to reproduce the figures we present here. The first vignette reproduces Figures 1 and 2 with an added explanation of the link between interaction coefficients and stochastic stability. The second vignette walks readers to our two step parameter estimation process, including the MAR CLS calculations, the stochastic stability calculations and classification of each community after a PCA. The third vignette gives an extended explanation of our RPM methodology as well as the code to reproduce Figures 7 and 8 as well as their equivalents for every single one of our data sets. The link to these vignettes is given in our Github readme file here https://github.com/jmponciano/StochasticMicrobiome/tree/main/ExampleCalcs.

## Data availability

Data are provisionally freely available at github.com/jmponciano and upon acceptance will be uploaded as well to the Dryad data repository and the Zenodo code repository.

## Code availability

Code is provisionally freely available at github.com/jmponciano and upon acceptance will be uploaded as well to the Dryad data repository and the Zenodo code repository.

## Acknowledgments

We would like to thank the valuable input and advice at various stages of this work from Drs Mark L Taper, Brian Dennis, and Robert D. Holt, C George Glen and Michelle L. Gaynor. Funding for this research was provided by the National Institute of General Medical Sciences of the National Institutes of Health, under grant number 1R01GM103604 to University of Florida, with JMP as PI. JMP was also supported by NSF grant No. 2052372 to University of Florida. This work was also in part supported by the National Institute of Allergy and Infectious Diseases Extramural Activities grant no. R01 AI084918 from the National Institutes of Health (NIH)

## Author contributions

J.M.P., L.J.F. and J.R. conceived the study. L.J.F. and J.R. are responsible for the molecular biology experiments and data preparation. J.M.P. and J.P.G. curated the output data and analyzed the data. J.M.P. conceived and wrote the data analyses, J.M.P. and J.P.G. conducted all computer programming analyzing the data. J.M.P. and L.J.F. co-wrote the manuscript, all authors contributed to revisions. All authors read and approved the final manuscript. J.M.P. and L.J.F. are “co-first author”.

