## Supplemental Material for "Inferring stability and persistence in the vaginal microbiome: A stochastic model of ecological dynamics"

#### Clinical Study Methods

Participants (N=135) were recruited in Birmingham, Alabama, United States of America, from the Personal Health Clinic, the Jefferson County Department of Health STD Clinic, as well as through advertisements in newspapers (Ravel et al. 2013). All participants in this study were broadly consented following guidelines of the Human Microbiome Project of the National Institutes of Health. Women between 18-45y were enrolled in the study and their ethnicities were 62% African American, 32% White, 5% Hispanic and 1% Asian. Women were excluded from the study if they were pregnant, used the NuvaRing® for contraception, were less than 6 months postpartum or breastfeeding, had chronic illnesses such as kidney failure, diabetes, or HIV/AIDS, were diagnosed with an STI at enrollment, or used systemic or intravaginal antibiotics or antimycotics in the 30 days prior to enrollment. At the time of enrollment, a research nurse administered sensitive questionnaires. These were used to gather information on socioeconomic and demographic factors, female hygiene practices and health behaviors, gynecological and obstetrical history, sexual history and practices, sexually transmitted disease history, date of last menstrual period, methods of birth control currently used, alcohol and drug use, and fitness status and practices.

At the baseline visit the research nurse also assessed pelvic symptoms, performed a limited physical examination, collected biological specimens (see below), and recorded any physical findings including vaginal discharge and easily induced bleeding, and assessed the occurrence of ectopy, edema, inflammation, or ulcerations. During a pelvic examination, the nurse collected materials for the clinical assessment of BV using the Amsel 1 and Nugent criteria 2. In addition, the nurse tested for vulvovaginal candidiasis by microscopy and collected swabs that were used to test for Trichomonas vaginalis, Neisseria gonorrhoeae and Chlamydia trachomatis using molecular and microbiological methods. Finally, serum was collected and subsequently tested for syphilis, herpes simplex virus (HVS) type 1/2 and HIV. Positive results from any of these tests resulted in exclusion from the study. Participants were also provided detailed instructions on sample collection and storage as well as information on preparing vaginal smears.

At the baseline visit participants were given the materials needed to collect samples for one week. They were also provided detailed instructions on procedures to be used for the self-collection of vaginal swabs, preparation of vaginal smears, and instructions for swab storage and transport back to the clinic. Daily each subject self-collected three mid-vaginal swabs: the first Copan E-Swab was used to prepare a smear that was later Gram stained and used to determine Nugent scores. This swab was then placed in Liquid Amies Transport Media and used later used for extracting genomic DNA. In addition, subjects measured vaginal pH using the

CarePlan® VpH test glove (Inverness Medical). Finally, a diary was completed each day using a standardized form on which all responses were pre-coded to record hygiene practices and sexual activities. These included information on the use of sanitary napkins, tampons, and douching, as well as vaginal intercourse, receptive oral sex, digital penetration, rectal sex, sex toys or the use of diaphragms, condoms, spermicides, lubricants. Women also reported menstrual bleeding, and vaginal symptoms that included vaginal itching, discharge, odor, irritation, and pain on urination. After collection, all samples were stored in the participants' home freezers. Each week the subjects transported their samples in a cooler to the study site where they were then transferred to a -80°C freezer. At this time another one-week sampling kit was provided to the study subjects. At weeks 5 and 10, the participants completed another detailed questionnaire, and had a thorough medical evaluation that included scoring for bacterial vaginosis using Amsel and Nugent criteria. Antibiotic treatment was offered to the participants if the conditions warranted.

All vaginal smears from daily sampling were Gram-stained and scored using Nugent criteria by personnel in Dr. Schwebke's laboratory at the University of Alabama. Over 9,000 slides were scored. In addition, batches of samples were shipped on dry ice to the Institute for Genome Sciences at the University of Maryland School of Medicine at weekly intervals whereupon the samples were again stored at -80°C. In total over 33,000 biological samples were collected in this study. All data from this study are managed and stored at the Institute for Genomic Research at the University of Maryland School of Medicine in a secure relational database that includes all de-identified metadata (medical evaluations, answers to all questionnaires, and daily diaries) and a system to track barcoded samples from each participant.

We have demonstrated that the long-term storage of samples at -80°C does not alter the vaginal microbiome and metabolome when compared to fresh samples (Bai et al. 2012). In a previous study we demonstrated that there were minimal differences between contemporaneously self-collected and physician-collected swabs samples collected from the same individual (Forney et al. 2010) as judged by the composition of vaginal communities determined by sequencing bacterial 16S rRNA genes. Others have reported similar findings (Menar et al. 2012, Nelson et al. 2003). Finally, all our methodology for DNA extraction, 16S rRNA gene amplification and sequencing and taxonomic assignments was published in Ravel et al 2013. All the data analyzed here is publicly avaible at NCBI's short read archive Bioproject number PRJNA208535.

#### PCA on stability metrics and stability classification

In the main text, Figure 5, we show a classification scheme of women according to the stability metrics estimated from fitting a MAR model to their bacterial time series data. The stability metrics for each woman computed from the parameter estimates of the two-species MAR model (Lactobacillus versus the rest) are shown in Supplementary Table 2. These stability metrics were then used to run a PCA with the observations being each woman and the variables being the four stability metrics presented in this table. The full code for the PCA was done in R following Johnson and Wichern (2002), chapter 8 and modified from JMP's statistics multivariate statistics teaching material. It is available at github.com/jmponciano so that all figures are reproducible. In Supplementary Table 3 we printed the correlation of each one of the four stability metrics with each principal component. The first three stability metrics (the variance proportion, the mean return time and the variance in the return time) have the highest negative correlation with the first principal component (PC I). The smaller the values in these three statistics, the more stable the dynamics is and the highest the PC I score. In this case, PC I explain 60.34% of the variability. PC II explains an additional 24.48% of the variance, so that together, the first two principal components explain 84.82% of the variation. The fourth stability metric, reactivity, has the highest (negative) correlation with PC II. Although devising a classification scheme based on these stability metrics can be achieved in multiple ways, basing some scheme on an ecological-processes rationale gives intuitive results. For example, in the PCA plotted on the main text, we colored the different women according to a qualitative stability scale, going from "very unstable" to "very stable". To derive such scale using ecological principles, we used each woman's score in the first two principal components scores as well as their overall PCA score and the mean strength of density dependence (the mean of the diagonal of the B matrix) of their bacterial communities as clustering variables in a k-means cluster. We set k=4. As with any cluster analysis, many different variables can be used to obtain a clustering/grouping scheme and the following is but one of the possible ways of achieving such grouping.

The cluster means are shown in Supplementary Table 4. Women with the highest score in PC I, which were the women with the lowest (on average) first three stability metrics and hence the women with the highest stability consistently appeared grouped in cluster 1. Those women also have on average the lowest mean density-dependent coefficient. Recall that the smaller that coefficient, the stronger the self-regulation (intra-specific density dependence) which according to Ives et al (2003) is also consistent with a more stable stochastic population dynamics. Hence, we classified the bacterial dynamics in these women as “highly stable”. Women in cluster 3 had on average the next highest score in PC I and the second smallest (on average) strength of density dependence. Hence, we classified the bacterial population dynamics in these women as “stable”. The dynamics of the bacterial communities in women on cluster 2 had the second highest average density-dependent coefficient, nearing the value of 1, which represents unregulated (density-independent) growth. The bacterial communities of these women also had an average PC I score that ranked third, following that of clusters 1 and 2, which means that the first three stability metrics estimates are higher than the rest, hence less stable. Finally, the communities in cluster 4 had PC scores that were the lowest on average and the highest mean density-dependence coefficient which neared 1 (0.98, see supplementary table 4). Hence, we labeled these communities as highly unstable. All analyses and documentation can be found in the R programs in github.com/jmponciano.

### Supplementary Figures

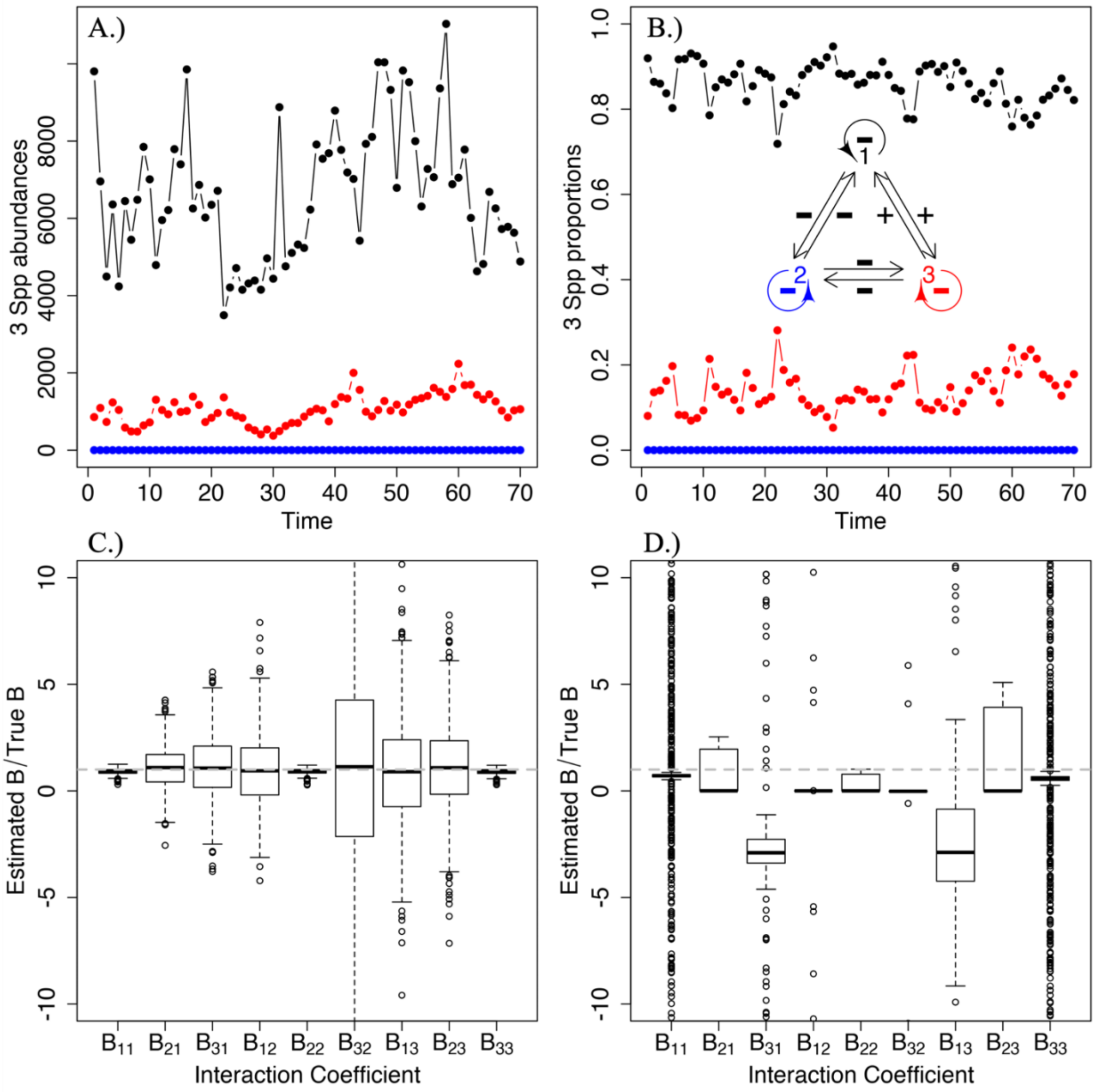


**Supplementary Figure 1:** Simulated populations trajectory during 70 days for a three-species community, and estimates of the interaction strengths. Panels A.) and B.) show the abundances on the left and the relative abundances on the right for the same simulation. Inset on B.) is a diagram representing the structure of the community using one color per species as in the plots. In this particular simulation setting, all the interactions were weak. The parameter values for the simulation are shown in Supplementary Table 1. Panels C.) and D.) show the boxplots of the relative bias of the estimates of all the interaction strengths between all species (the $B_{ij}$, $i = 1,2,3$) obtained using the total abundances on the left and the relative abundances on the right. To do these boxplots, 1000 simulations under this particular community structure and parameter values were done. Boxplots centered around the dotted gray line at 1 denote unbiased estimates. See text for details.

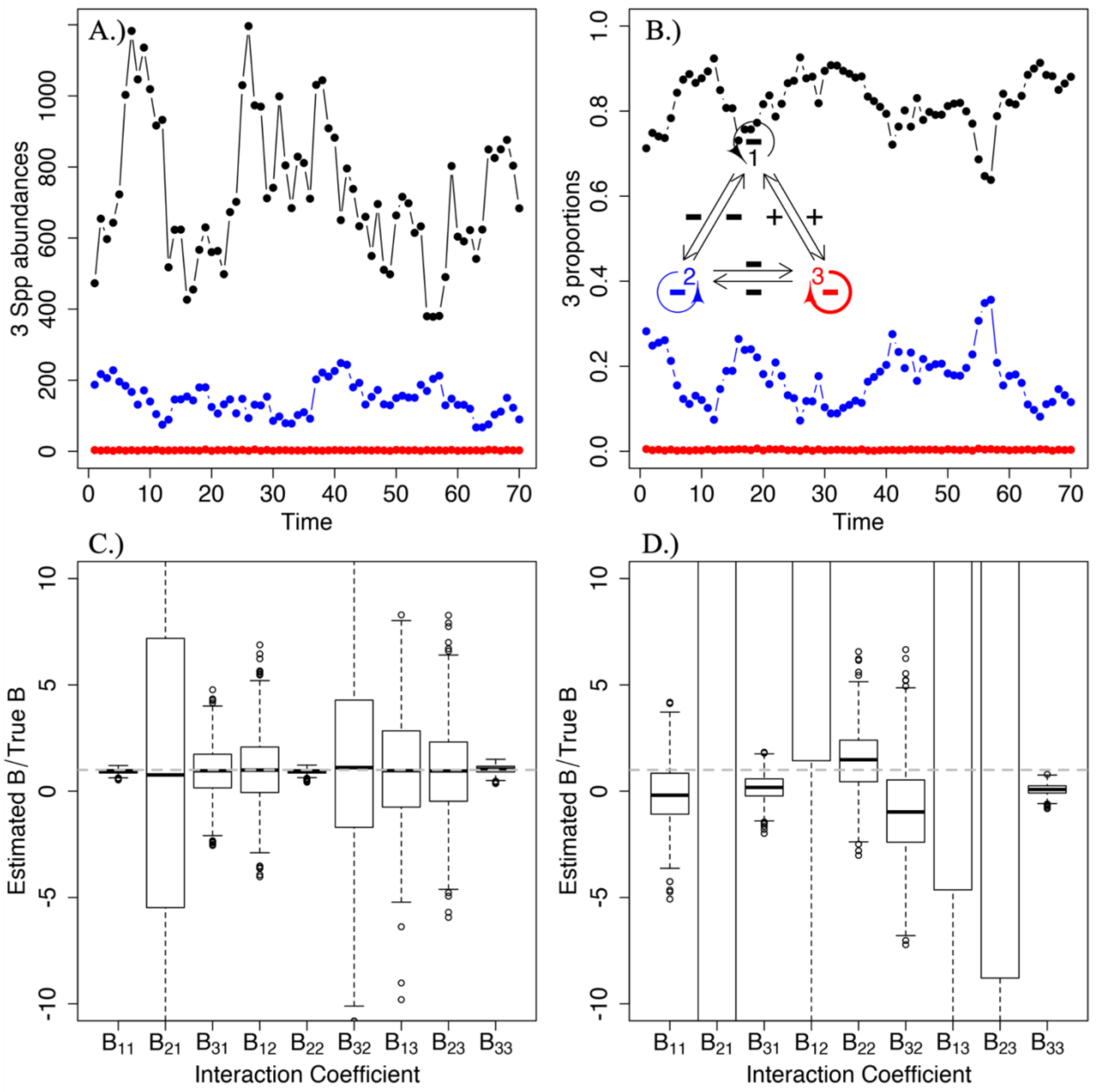


Supplementary Figure 2: Simulated populations trajectory during 70 days for a three-species community, and estimates of the interaction strengths. Panels A.) and B.) show the abundances on the left and the relative abundances on the right for the same simulation. Inset on B.) is a diagram representing the structure of the community using one color per species as in the plots. In this particular simulation setting, all the interactions were weak except for the strength of intra-specific competition, or density dependence, for species 3. The parameter values for the simulation are shown in Supplementary Table 1. Panels C.) and D.) show the boxplots of the relative bias of the estimates of all the interaction strengths between all species (the $B_{ij}$, $i = 1,2,3$) obtained using the total abundances on the left and the relative abundances on the right. To do these boxplots, 1000 simulations under this particular community structure and parameter values were done. Boxplots centered around the dotted gray line at 1 denote unbiased estimates. See text for details.

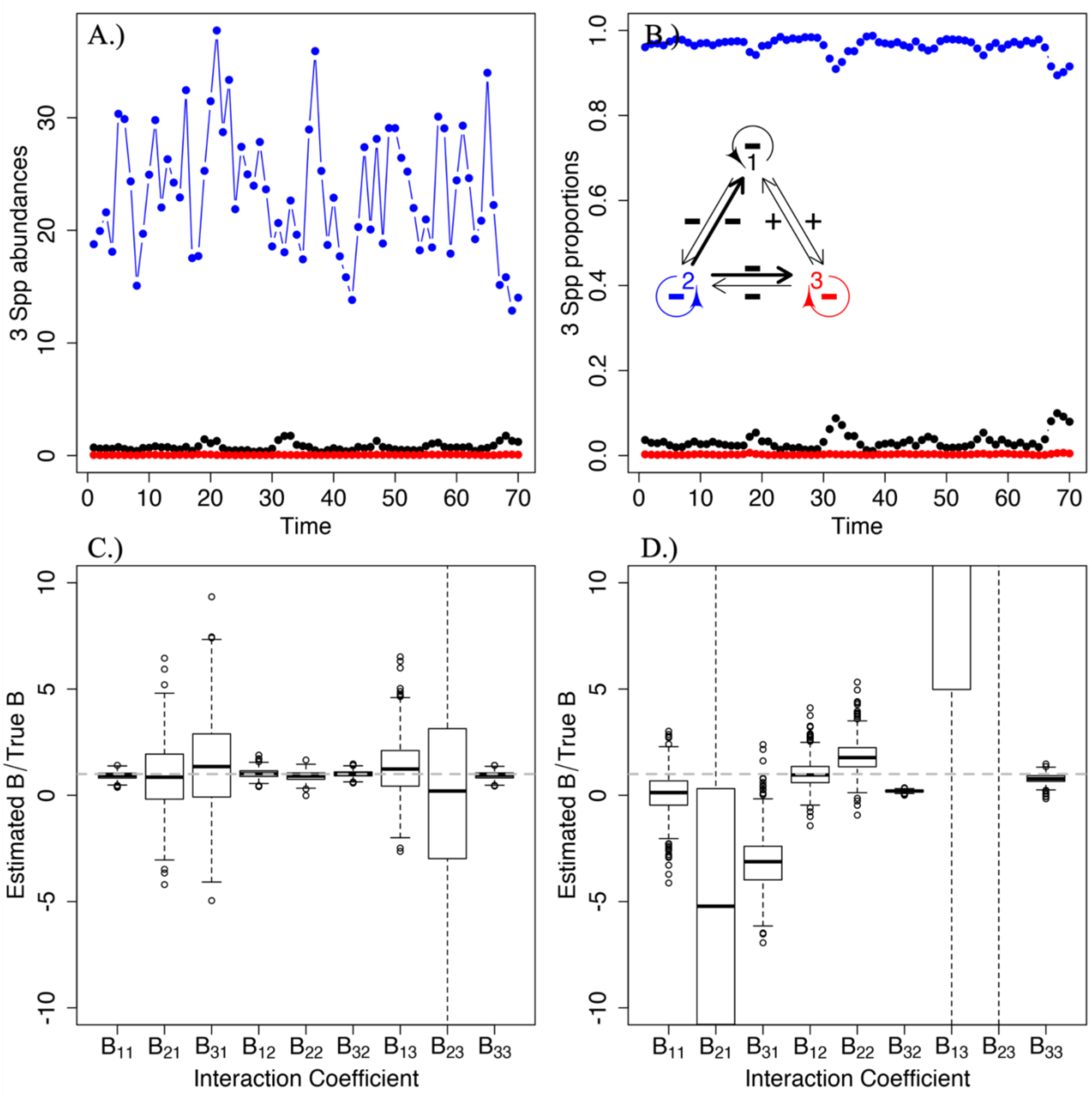


Supplementary Figure 3: Simulated populations trajectory during 70 days for a three-species community, and estimates of the interaction strengths. Panels A.) and B.) show the abundances on the left and the relative abundances on the right for the same simulation. Inset on B.) is a diagram representing the structure of the community using one color per species as in the plots. In this particular simulation setting, all the interactions were weak except for the strength of inter-specific competition, from species 2 to species 3 and 1. The parameter values for the simulation are shown in Supplementary Table 1. Panels C.) and D.) show the boxplots of the relative bias of the estimates of all the interaction strengths between all species (the $B_{ij}$, $i = 1,2,3$) obtained using the total abundances on the left and the relative abundances on the right. To do these boxplots, 1000 simulations under this particular community structure and parameter values were done. Boxplots centered around the dotted gray line at 1 denote unbiased estimates. See text for details.

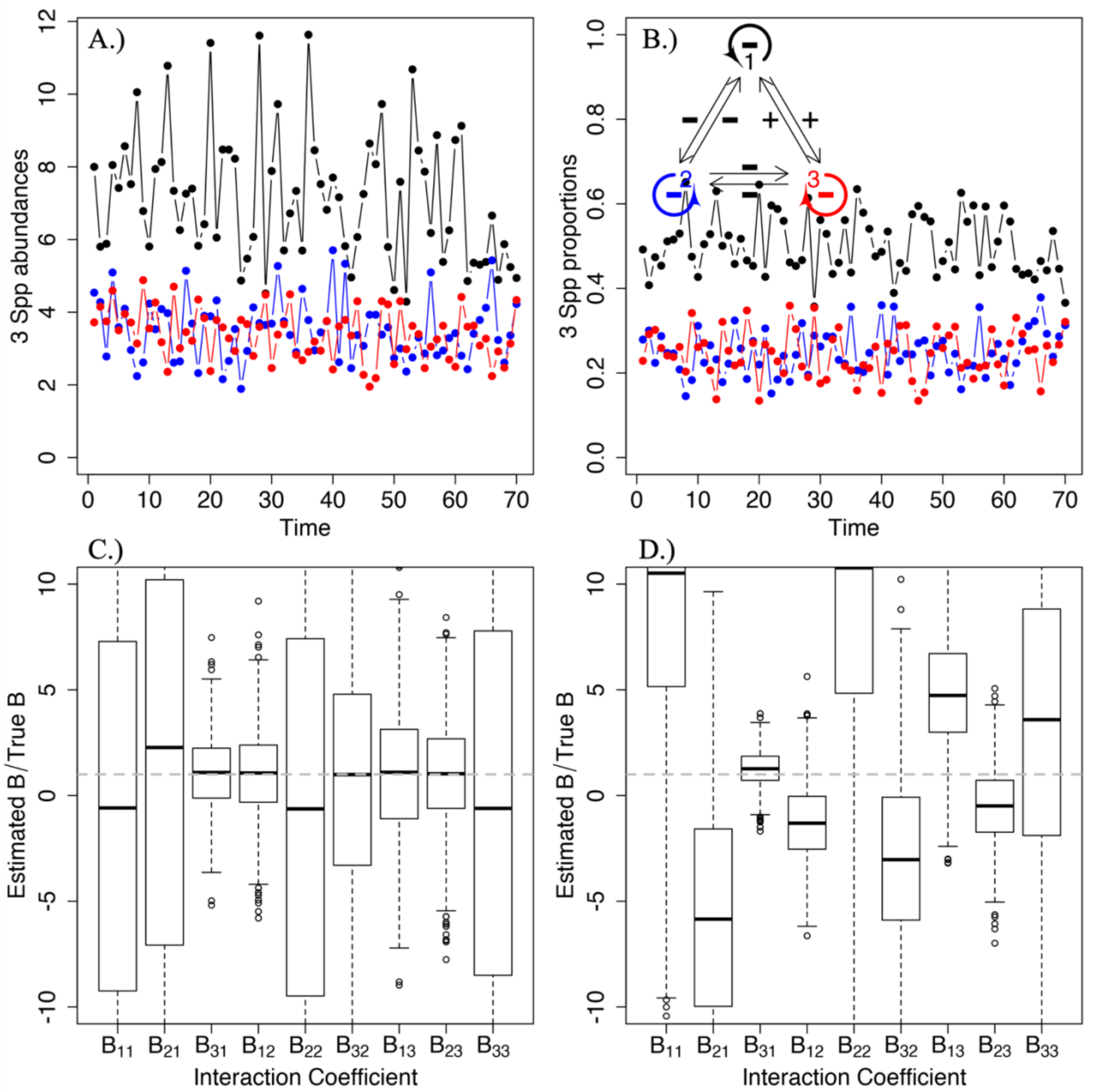


Supplementary Figure 4: Simulated populations trajectory during 70 days for a three-species community, and estimates of the interaction strengths. Panels A.) and B.) show the abundances on the left and the relative abundances on the right for the same simulation. Inset on B.) is a diagram representing the structure of the community using one color per species as in the plots. In this particular simulation setting, all inter-specific interactions were weak and all intra-specific interactions, or density dependence values, were strong. The parameter values for the simulation are shown in Supplementary Table 1. Panels C.) and D.) show the boxplots of the relative bias of the estimates of all the interaction strengths between all species (the $B_{ij}$, $i = 1,2,3$) obtained using the total abundances on the left and the relative abundances on the right. To do these boxplots, 1000 simulations under this particular community structure and parameter values were done. Boxplots centered around the dotted gray line at 1 denote unbiased estimates. See text for details.

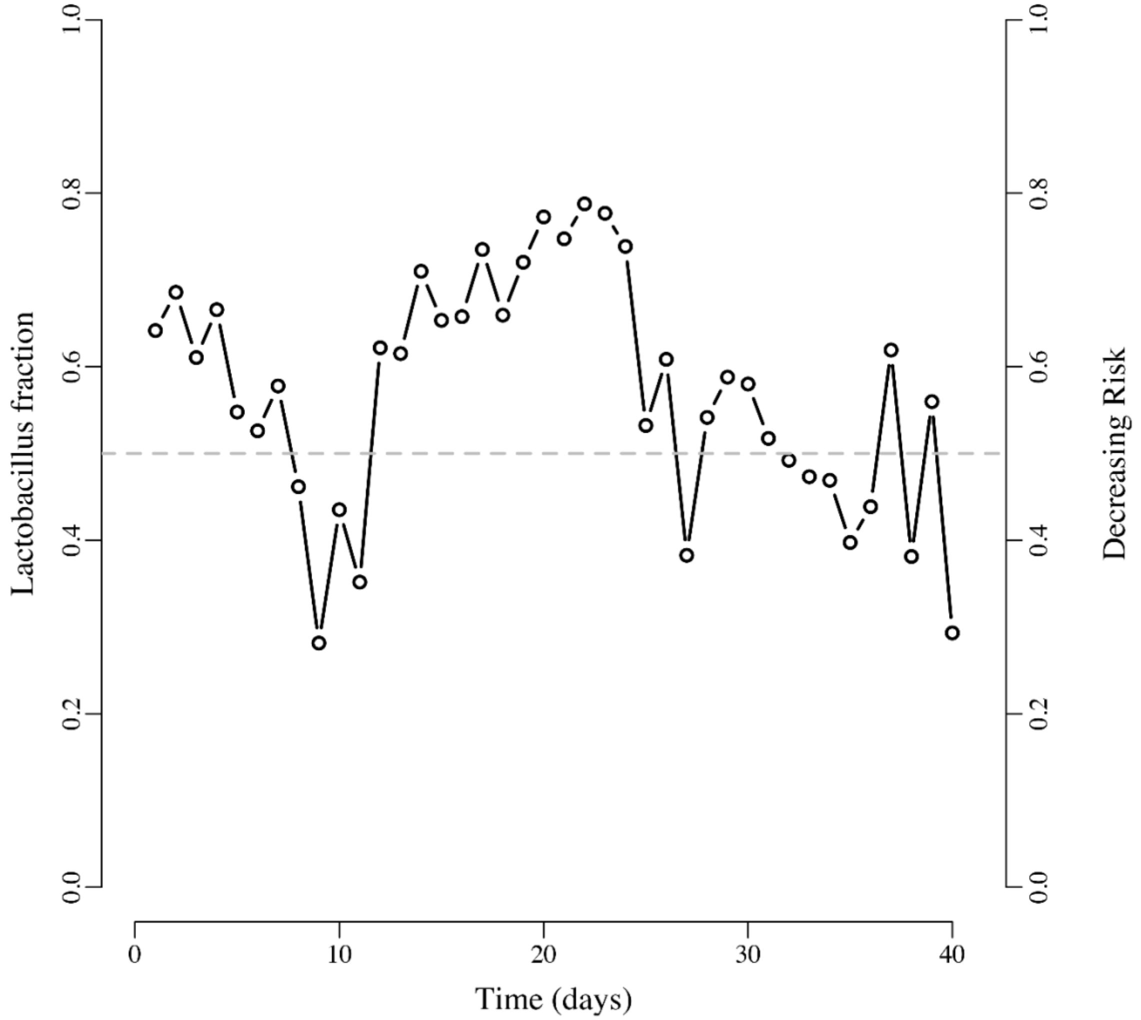


**Supplementary Figure 5:** When the relative abundance of *Lactobacillus* dwindles down below a 0.5 proportion, the bacterial community is under a high risk of infection by HIV (Klatt et al 2017). On the other hand, as the relative abundance of *Lactobacillus* moves above 0.5, the risk of infection decreases. Seeking to elucidate which type and magnitude of ecological interactions would lead to desirable dynamics (i.e. fluctuations in relative abundance of *Lactobacillus* above 0.5) is a reachable target under our analysis using the MAR model.

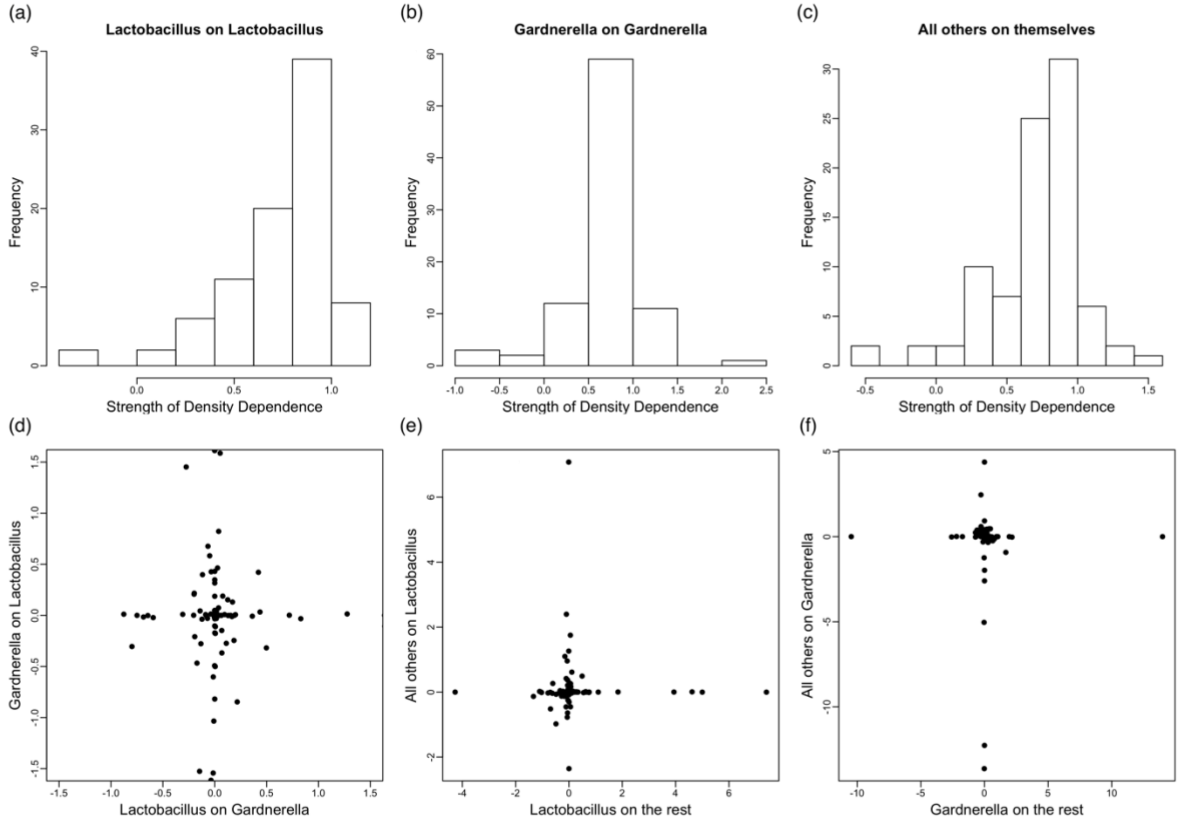


**Supplementary Figure 6.** Variability across women of the interaction relationships between three groups of species. This figure illustrates the wide variability of interaction coefficients within the same pair of species for our three-species model fit, where all *Lactobacillus* were grouped together, *Gardnerella* was kept as a separate second species and all the other species as a third functional group. Take for instance the two-way interaction strengths between *Gardnerella* and *Lactobacillus*. Across all 88 women, one sees interaction strengths in all quadrants: +/+, +/-, -/+ and -/-.

#### Supplementary Tables

**Supplementary Table 1.** Parameters used in the simulation of the three species community time series based on four different scenarios (see main text, Figure 2 and Supplementary Figures 1-4). The first row corresponds to the vector of maximum growth rates for every species. The next three rows correspond to the values used for the variance-covariance matrix of the environmental variation and finally, the next four sets of three rows correspond each to the matrix of interaction coefficients B. The (i,j) element in these 3 by 3 tables correspond to the effect of species j on the growth rate of species i.

| | Species 1 | Species 2 | Species 3 |
|---|---|---|---|
| *A* | 1.9 | 1.3 | 1.1 |
| $\Sigma$ | | | |
| Species 1 | 0.05 | 0.005 | 0.005 |
| Species 2 | 0.005 | 0.05 | 0.005 |
| Species 3 | 0.005 | 0.005 | 0.05 |
| matrix, scenario 1 | | | |
| Species 1 | 0.75 | -0.06 | 0.04 |
| Species 2 | -0.1 | 0.75 | -0.05 |
| Species 3 | 0.07 | -0.02 | 0.75 |
| *B* matrix, scenario 2 | | | |
| Species 1 | 0.75 | -0.06 | 0.04 |
| Species 2 | -0.01 | 0.75 | -0.05 |
| Species 3 | 0.07 | -0.02 | -0.5 |
| *B* matrix, scenario 3 | | | |
| Species 1 | 0.55 | -0.60 | 0.07 |
| Species 2 | -0.06 | 0.55 | -0.02 |
| Species 3 | 0.04 | -0.75 | 0.55 |
| *B* matrix, scenario 4 | | | |
| Species 1 | 0.01 | -0.06 | 0.04 |
| Species 2 | -0.01 | 0.01 | -0.05 |
| Species 3 | 0.07 | -0.02 | 0.01 |

Supplementary Table 2. Stability metrics for each woman computed from the parameter estimates of the two-species MAR model (Lactobacillus versus the rest)

| Individual | Variance Proportion | Mean Return Time | Variance Return Time | Reactivity |
|---|---|---|---|---|
| woman1 | 0.610 | 0.973 | 0.946 | -0.018 |
| woman2 | 0.676 | 0.962 | 0.926 | -0.034 |
| woman3 | 0.167 | 0.954 | 0.910 | -0.719 |
| woman4 | 0.127 | 0.919 | 0.845 | -0.077 |
| woman5 | 0.767 | 0.998 | 0.996 | -0.561 |
| woman6 | 0.134 | 0.997 | 0.993 | -0.064 |
| woman7 | 0.920 | 1.004 | 1.009 | 2.793 |
| woman8 | 0.739 | 0.927 | 0.860 | -0.115 |
| woman10 | 0.598 | 0.879 | 0.773 | -0.147 |
| woman11 | 0.842 | 0.997 | 0.993 | -7.613 |
| woman13 | 0.691 | 0.989 | 0.978 | -0.016 |
| woman14 | 0.012 | 0.782 | 0.612 | -0.011 |
| woman15 | 0.307 | 0.935 | 0.875 | -0.215 |
| woman16 | 0.332 | 0.897 | 0.805 | -0.039 |
| woman17 | 0.346 | 0.992 | 0.983 | -0.184 |
| woman18 | 0.607 | 0.995 | 0.989 | -16.015 |
| woman19 | 0.569 | 0.998 | 0.996 | -36.740 |
| woman21 | 0.700 | 0.995 | 0.991 | -2.040 |
| woman22 | 0.910 | 0.994 | 0.988 | -6.360 |
| woman23 | 0.461 | 0.965 | 0.932 | -0.160 |
| woman26 | 0.480 | 0.968 | 0.937 | -0.300 |
| woman27 | 0.046 | 0.974 | 0.948 | -0.951 |
| woman28 | 0.501 | 0.998 | 0.996 | -20.550 |
| woman29 | 0.541 | 0.949 | 0.901 | -0.382 |
| woman30 | 0.001 | 0.348 | 0.121 | -2.881 |
| woman31 | 0.007 | 0.588 | 0.346 | -0.034 |
| woman35 | 0.534 | 0.957 | 0.916 | -0.842 |
| woman36 | 0.153 | 0.974 | 0.948 | -0.002 |
| woman38 | 0.053 | 0.995 | 0.990 | -3.389 |
| woman39 | 0.555 | 0.996 | 0.991 | -184.750 |
| woman41 | 0.657 | 0.997 | 0.995 | -0.159 |
| woman42 | 0.530 | 0.905 | 0.819 | -0.030 |
| woman43 | 0.486 | 0.986 | 0.971 | -0.002 |
| woman44 | 0.232 | 0.870 | 0.757 | -0.063 |
| woman46 | 0.410 | 0.946 | 0.895 | -1.742 |
| woman47 | 0.291 | 0.752 | 0.566 | -0.040 |
| woman48 | 0.624 | 0.967 | 0.935 | -0.926 |
| woman49 | 0.718 | 0.940 | 0.884 | -0.427 |
| woman50 | 0.002 | 0.753 | 0.567 | -0.079 |
| woman52 | 0.197 | 0.906 | 0.820 | -0.167 |
| woman53 | 0.761 | 0.932 | 0.872 | -0.488 |

| | | | | |
|---|---|---|---|---|
| woman55 | 0.369 | 0.945 | 0.893 | -0.048 |
| woman56 | 0.116 | 0.886 | 0.784 | -0.006 |
| woman58 | 0.560 | 0.925 | 0.855 | -0.008 |
| woman59 | 0.513 | 0.832 | 0.716 | -0.141 |
| woman60 | 0.421 | 0.847 | 0.718 | -0.087 |
| woman61 | 0.305 | 0.921 | 0.847 | -0.452 |
| woman62 | 0.414 | 0.986 | 0.971 | -0.002 |
| woman65 | 0.001 | 0.865 | 0.748 | -2.620 |
| woman66 | 0.084 | 0.984 | 0.968 | -0.019 |
| woman69 | 0.290 | 0.819 | 0.672 | -0.181 |
| woman70 | 0.510 | 0.925 | 0.856 | -0.230 |
| woman71 | 0.402 | 0.962 | 0.926 | -0.890 |
| woman75 | 0.499 | 0.908 | 0.824 | -0.062 |
| woman76 | 0.523 | 0.917 | 0.841 | -0.020 |
| woman77 | 0.812 | 0.949 | 0.901 | -0.042 |
| woman79 | 0.741 | 1.003 | 1.006 | 0.023 |
| woman82 | 0.446 | 0.989 | 0.977 | -0.046 |
| woman83 | 0.107 | 0.762 | 0.580 | -0.110 |
| woman87 | 0.686 | 0.997 | 0.993 | -0.232 |
| woman88 | 0.142 | 0.797 | 0.635 | -0.074 |
| woman90 | 0.848 | 1.010 | 1.021 | -0.008 |
| woman92 | 0.478 | 0.936 | 0.877 | -0.012 |
| woman93 | 0.330 | 0.758 | 0.574 | -0.069 |
| woman96 | 0.759 | 0.959 | 0.919 | -0.026 |
| woman97 | 0.609 | 0.928 | 0.862 | -0.053 |
| woman101 | 0.001 | 0.934 | 0.872 | -0.026 |
| woman102 | 0.015 | 0.983 | 0.966 | -0.031 |
| woman103 | 0.325 | 0.899 | 0.809 | -0.034 |
| woman112 | 0.527 | 0.932 | 0.869 | -0.662 |
| woman114 | 0.126 | 0.935 | 0.874 | -0.218 |
| woman115 | 0.136 | 0.794 | 0.630 | -4.959 |
| woman116 | 0.239 | 0.957 | 0.915 | -0.868 |
| woman117 | 0.808 | 0.996 | 0.993 | -0.082 |
| woman118 | 0.556 | 0.928 | 0.862 | -0.118 |
| woman119 | 0.307 | 0.971 | 0.942 | -0.025 |
| woman120 | 0.486 | 0.961 | 0.923 | -0.006 |
| woman121 | 0.572 | 0.960 | 0.921 | -0.105 |
| woman122 | 0.329 | 0.996 | 0.992 | -5.526 |
| woman124 | 0.692 | 0.977 | 0.954 | -0.028 |
| woman125 | 0.811 | 0.997 | 0.994 | -0.188 |
| woman126 | 0.001 | 0.828 | 0.686 | -0.020 |
| woman128 | 0.711 | 0.998 | 0.996 | -0.793 |
| woman129 | 0.522 | 0.850 | 0.723 | -0.010 |
| woman130 | 0.744 | 0.970 | 0.941 | -0.303 |
| woman131 | 0.098 | 0.792 | 0.626 | -0.171 |
| woman134 | 0.029 | 0.894 | 0.800 | -1.005 |

| | | | | |
|---|---|---|---|---|
| woman135 | 0.793 | 1.000 | 1.000 | -0.006 |

Supplementary Table 3. Correlation of each one of the four stability metrics with each principal component. The data for the PCA is shown in Supplementary Table 2 above.

| Variable | PC1 | PC2 | PC3 | PC4 |
|---|---|---|---|---|
| Variance Proportion | -0.729 | -0.096 | 0.678 | 0.003 |
| Mean Return time | -0.956 | -0.068 | -0.278 | 0.071 |
| Variance Return time | -0.966 | -0.054 | -0.243 | -0.073 |
| Reactivity | 0.190 | -0.981 | -0.034 | -0.001 |

Supplementary Table 4. Centroids from each cluster resulting from a k-means cluster (k=4) of the 88 women with four variables. The variables were: the PC I and II scores, the overall PCA standarized score (the eigenvector times the standarized values, eq. 8-29 Johnson and Wichern (2002)) and the average density-dependent coefficient in the bacterial community (the average of the diagonal entries in the **B** matrix of the MAR model). As with any cluster analysis, many different variables can be used to obtain a clustering/grouping scheme and the following is but one of the possible ways of achieving such grouping.

| Clusters | Scaled Scores 1 | Scaled Scores 2 | PCA scores | Mean ddp |
|---|---|---|---|---|
| 1 | 0.223 | 0.027 | 2.071 | 0.521 |
| 2 | -0.016 | -0.018 | -0.226 | 0.786 |
| 3 | 0.056 | 0.057 | 0.586 | 0.690 |
| 4 | -0.090 | -0.020 | -0.780 | 0.906 |
